# The vulnerability of overwintering insects to loss of the subnivium

**DOI:** 10.1101/2024.05.06.592805

**Authors:** Kimberly L. Thompson, Jonathan N. Pauli, Benjamin Zuckerberg

## Abstract

Winter climate change threatens the subnivium (i.e., the microhabitat that exists between the snowpack and the ground), and the community of species that depends on it for overwintering survival. One group of species that will likely exhibit an array of responses to subnivium loss are overwintering insects because they vary in their cold tolerance strategies and lower thermal limits. For an assemblage of eight insect species that range in their cold tolerance strategies and include both pollinators and pests, we investigated species-specific vulnerabilities to shifting subnivium conditions by applying information on each species’ supercooling point to spatially- and temporally-explicit models of minimum subnivium temperatures for three warming scenarios in the Great Lakes region in the United States: current conditions (i.e., control), +3°C, and +5°C. Although species varied in their vulnerabilities, our predictions indicated that exposure to lethal temperatures generally decreased under warming of 3°C, but increased under warming of 5°C, indicating that once enough warming happens, a tipping point is reached. We also found that freeze-tolerant species (i.e., species that can survive at temperatures below their supercooling point) possess a more cryptic vulnerability to winter climate change because sustained below-freezing temperatures were sufficient to induce vulnerability (i.e., predicted mortality), even when temperatures were above the supercooling point. This work provides a better understanding of the vulnerability of different insect species to winter climate change, which is critical because overwintering survival and the fitness consequences incurred during overwintering likely represent important bottlenecks for the population dynamics of subnivium-dependent species.

## Introduction

Global climate change disrupts a host of abiotic processes (Campbell, et al. 2009, Gao, et al. 2015) that together have altered the conditions in which species persist (Herrera, et al. 2018, Lehikoinen, et al. 2011). This is especially true for winter in temperate regions, where climate change has led to reductions in snow cover extent and duration (Lemke and Ren 2007, Serreze 2010), as well as an increase in extreme events (e.g., extreme cold outbreaks, polar vortices) (Kodra, et al. 2011, Vavrus, et al. 2006). Future climate scenarios predict continued reductions in snow cover due to the conversion of snowfall to rainfall and more frequent rain-on-snow events (Peacock 2012), which will result in shallower, denser, and more variable snowpacks (Christopher, et al. 2008). These changes give rise to one of the most critical consequences of winter climate change: the degradation of the subnivium, the microhabitat that exists between the snowpack and the ground (Pauli, et al. 2013, Thompson, et al. 2021).

Since the low thermal conductivity of snow traps heat released from the soil (Pruitt 2005), the subnivium provides a thermally stable refuge for a diversity of overwintering plants and animals (Pauli, et al. 2013). Therefore, sufficient depths of low-density snow keep ground temperatures stably around 0°C regardless of fluctuations in ambient air temperature (Marchand 2013). Warmer winter temperatures, however, reduce the extent, duration, and thermal stability of the subnivium (Thompson, et al. 2018, Thompson, et al. 2021), leading to more variable ground temperatures and freeze-thaw cycles (Brown and DeGaetano 2011, Grillakis, et al. 2016, Groffman, et al. 2001). With reduced snow cover, ground temperatures are also more susceptible to prolonged cold from extreme cold outbreaks. Consequently, loss of the subnivium microhabitat will have severe ecological consequences for the community of species that depend on it for overwintering survival.

Although a compromised subnivium is more susceptible to colder and more variable winter temperatures that can reduce survival rates of species overwintering there (Bokhorst, et al. 2012, Korslund and Steen 2006, O’Connor and Rittenhouse 2016), the effects of subnivium loss will not be uniform across taxa, given the wide diversity of adaptations to cold stress. Despite a growing focus in recent years on merging species distribution modeling with metabolic requirements and microclimate conditions (Kearney and Porter 2009, Lenoir, et al. 2017), few studies have addressed overwintering species in the subnivium (but see Kearney 2020), and those that have tend to be single-species focused due to the amount of data and computation time required (e.g., Fitzpatrick, et al. 2019). Since the effects of warmer winter temperatures and subnivium loss on overwintering survival are likely to be complex, understanding the relative impacts on assemblages of overwintering species is necessary for directing future research and conservation efforts.

One group of species that will likely exhibit an array of responses to subnivium loss are overwintering insects because they have a wide range of cold tolerance strategies, including freeze tolerance, freeze avoidance, chill tolerance, and chill susceptibility (Lee 2010, Overgaard and MacMillan 2017). Freeze-tolerant species synthesize ice-nucleating agents in the winter, which enable conversion of up to 80% of their extracellular bodily fluids to ice at temperatures at or above -10°C (Brown, et al. 2004, Lee 2010). This extracellular freezing protects organs and tissues by preventing the irreversible damage that would result from ice crystal formation inside cells (Toxopeus and Sinclair 2018). The temperature at which extracellular freezing occurs is known as the supercooling point (SCP) (Lee 2010), i.e., the temperature at which insects can avoid ice formation in their cells and survive at sub-freezing temperatures (Dancau, et al. 2018). After temperatures fall below the SCP, freeze-tolerant individuals can survive additional cooling, but there is a high amount of interspecific variation in the difference between the SCP and the temperature at which mortality occurs, known as the lower lethal temperature (LLT) (Bale and Worland 2005). In cases where the LLT is close to, but still below the SCP, species are considered weakly freeze tolerant; when the LLT is far below the SCP, species are strongly freeze tolerant (Hart and Bale 1998, Sinclair, et al. 2015).

Contrary to freeze-tolerant species, freeze-avoidant species actively remove ice-nucleating agents from their bodies and produce cryoprotectants in their hemolymph that help them to remain in a liquid state at low temperatures without the formation of body ice (Brown, et al. 2004, Neven, et al. 1986). These cryoprotectants are effective up until the SCP, which in freeze-avoidant species is equivalent to the LLT. Temperature acclimation helps in these efforts, with brief exposure to nonlethal low temperatures triggering the accumulation of cryoprotectants in a process called rapid cold hardening (Kelty and Lee 1999). While freeze-avoidant species can survive cold temperatures as long as ice does not form in their bodies, for chill-susceptible species, mortality occurs at or around 0°C (Lee 2010, Sinclair, et al. 2015). Chill-tolerant species are somewhat more robust; they can withstand cold temperatures, but mortality occurs at temperatures above their SCP (Overgaard and MacMillan 2017).

Prior research on insect cold tolerance and overwintering success has typically assumed a constant organismal response to the environment (Marshall and Sinclair 2012) or a constant environmental input for the organism (Beekman, et al. 1998), yet both the environment and organismal responses to changes in the environment are highly variable. Additionally, although snow cover has been acknowledged as an important component of winter survival (Berzitis, et al. 2017, Brown, et al. 2004, Marshall and Sinclair 2012, Szabo and Pengelly 1973), many empirical studies have not used realistic values of subnivium temperatures, with experimental temperatures 3–4°C above the subnivium’s characteristically stable temperature of 0°C (Mercader and Scriber 2008, Scriber, et al. 2012, Woodard, et al. 2019). Consequently, there is a gap in knowledge about the physiological limits imposed by the environment and the relative impacts of climate change for different insect species that reside in the subnivium.

Here we unveil a framework to better understand the relative effects of winter climate change using an assemblage of insect species that range in their cold tolerance strategies, as well as the ecosystem services they provide. We apply information on the supercooling points for this set of species to models of current and future subnivium conditions in the Great Lakes region to better understand the vulnerability of these species in the face of winter climate change (Thompson, et al. 2021). By merging previously collected physiological data with a warming experiment that incorporates natural environmental variability and fine-scale drivers measured at a daily timescale and over a broad geographic extent, we quantify interspecific overwintering vulnerability across an entire winter season. Since the subnivium is predicted to be fairly resilient to warming of 3°C (Thompson, et al. 2021), we hypothesized that vulnerability across insect species would be roughly equivalent between current conditions and a warming scenario of 3°C, regardless of interspecific variation in supercooling points. Alternatively, since warming of 5°C is predicted to result in drastic reductions in subnivium extent and duration in all areas except those with lake-effect snow (Thompson, et al. 2021), we expected the vulnerability of insect species to follow a similar geographic pattern. Further, for species with lower SCPs, we hypothesized that the extent of their vulnerability would not be as widespread as those with higher SCPs.

## Methods

### Study area

The climate of the Great Lakes region is temperate, with warm summers and cold winters (ranging from approximately 8°C to 29°C and -22°C to 3°C, respectively) that vary according to latitude and proximity to the Great Lakes (Andresen, et al. 2014, PRISM Climate Group 2020). This area experiences the largest range of temperatures in the winter months, and the coldest overall temperatures occur in northern interior areas, away from the Great Lakes (Andresen, et al. 2014). Areas in the lake effect zone (i.e., downwind of the lakes) typically have more moderate climates with larger amounts of snowfall (Burnett, et al. 2003, Changnon and Jones 1972, Scott and Huff 1996).

### Species Selection

We surveyed the literature to find insect species that could be affected by changes in subnivium conditions. We selected species that: 1) have distributional ranges in all or part of the Great Lakes region; 2) overwinter in the subnivium or just under the soil surface (i.e., approximate subnivium); and 3) have published information on their supercooling points in the literature. Since the cold tolerance strategies of freeze tolerance and freeze avoidance are more common in temperate areas like the Great Lakes region (Overgaard and MacMillan 2017), we focused on species with these strategies, as well as a range of ecosystem services (e.g., pollinators and pests). Based on these criteria we selected eight species: the rusty patched bumblebee (*Bombus (Bombus) affinis*, freeze-avoidant pollinator), the yellow-banded bumblebee (*Bombus (Bombus) terricola*, freeze-avoidant pollinator), the diamondback moth (*Plutella xylostella*, freeze-avoidant pest), the Canadian tiger swallowtail (*Papilio canadensis*, freeze-avoidant pollinator), the Eastern tiger swallowtail (*Papilio glaucus*, freeze-avoidant pollinator), the woolly bear caterpillar (*Pyrrharctia Isabella*, freeze-tolerant pollinator), the bean leaf beetle (*Ceratoma trifurcata*, freeze-tolerant pest), and the hoverfly (*Syrphus ribesii*, freeze-tolerant pollinator). Further details on the characteristics of each species are supplied in the Supplementary Methods.

### Extraction of Species Data

Supercooling point data on bumblebees (*Bombus spp.*) and hoverflies (*Syrphus ribesii*) were extracted directly from published tables, while for all other species we used the software xyscan to extract data from published figures (Ullrich 2020). For the two bumblebee species (*B. affini*s and *B. terricola*), we used *Bombus terrestris*, a species in the same subgenus as *B. affinis* and *B. terricola* as a proxy due to the paucity of data on supercooling points in bumblebees (Supplementary Methods) (Cameron, et al. 2007). From the SCP values obtained for all species, we selected the lowest and highest reported SCP (highest/lowest mean ± SE/CI), to use in the analysis as a measure of the best- and worst-case scenario for each species (Table 1).

**Table 1.**
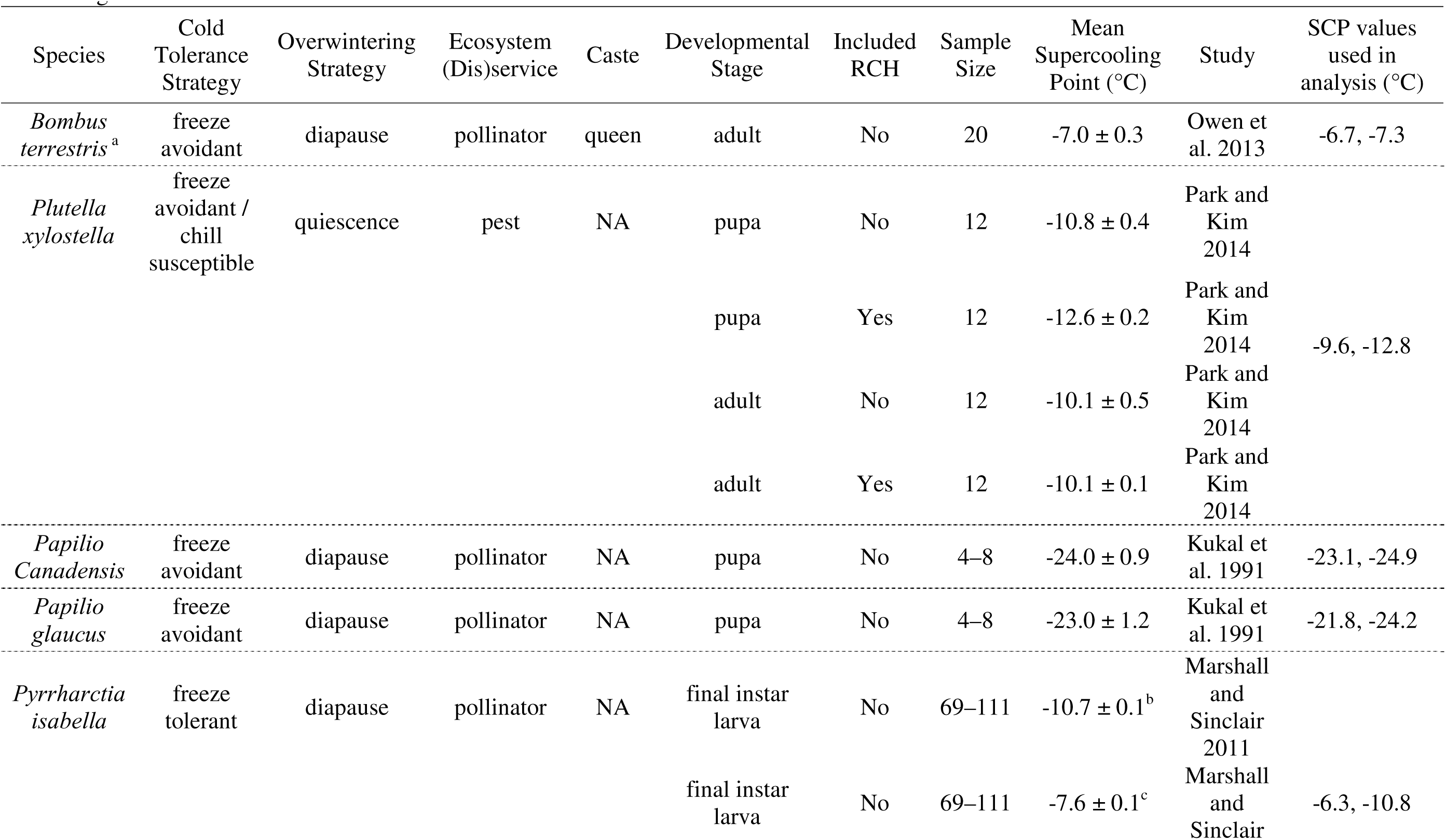
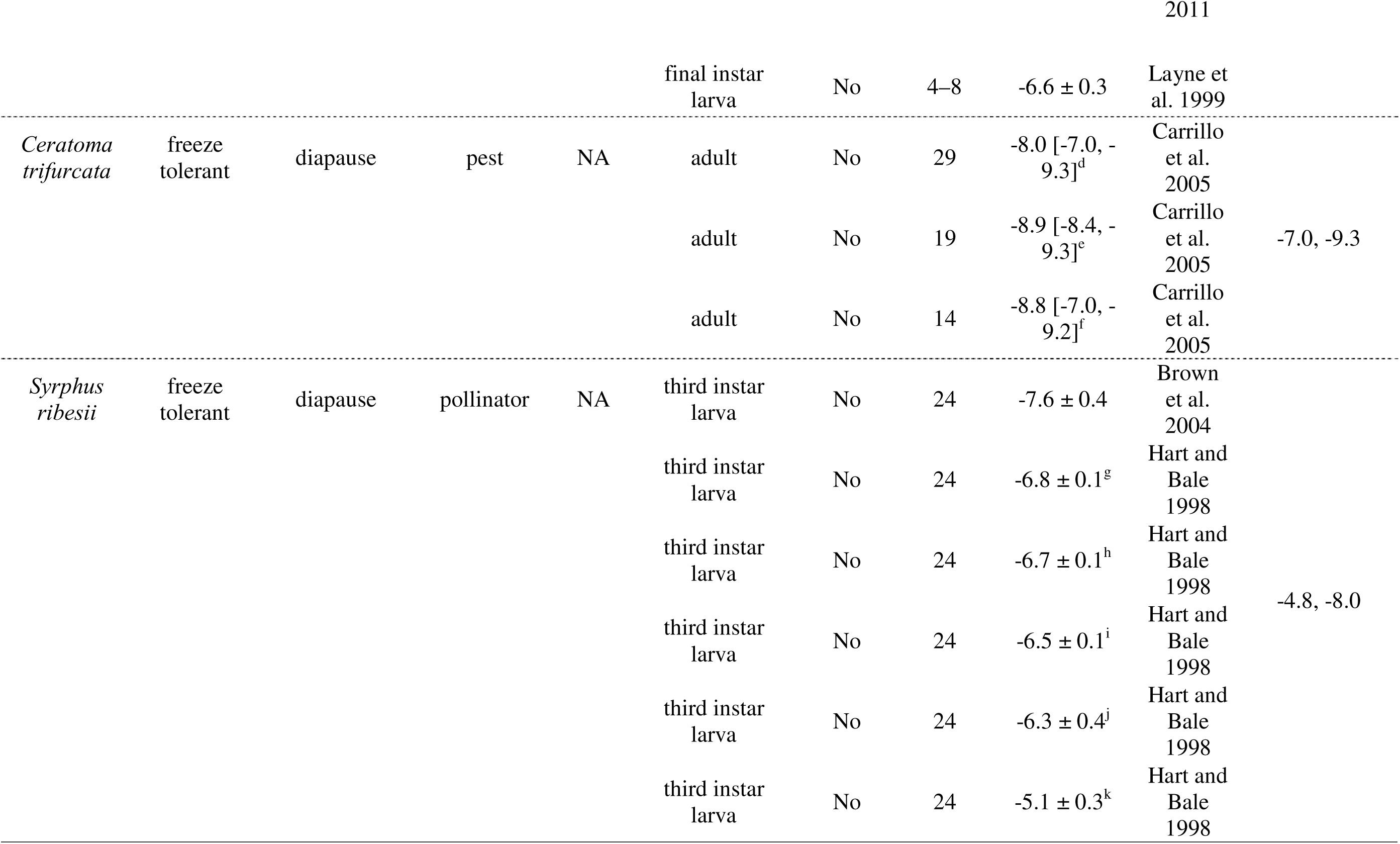
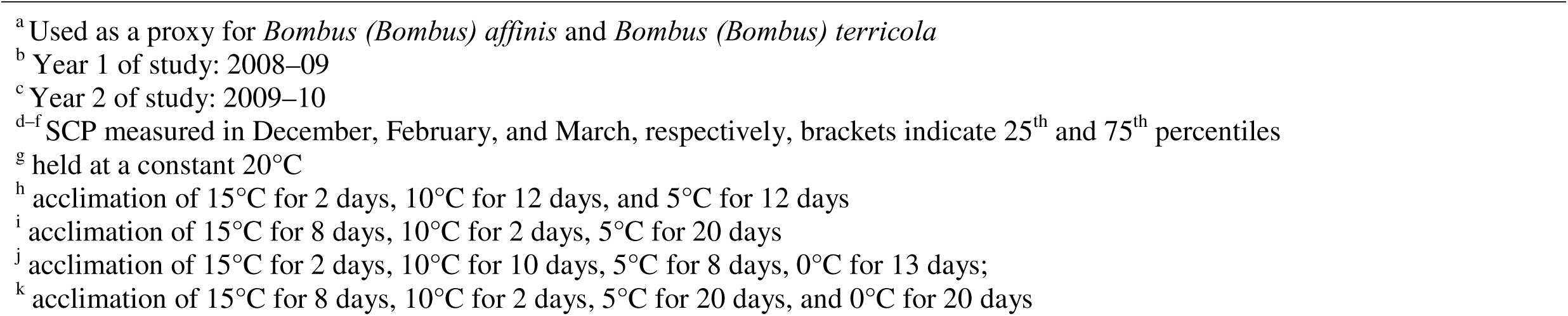
Species characteristics and supercooling points (SCP) extracted from published literature. We used the highest and lowest values (SCP ± Standard Error or Confidence Interval) in our analyses to represent a best and worst case for each species, respectively. Caste is specific to *Bombus spp.* and refers to the caste that overwinters in the subnivium. Developmental stage refers to the life stage (e.g., egg, larva, pupa, adult) during which overwintering occurs. Included RCH refers to whether each study included an acclimation period, otherwise known as rapid cold hardening.

For bean leaf beetles (*Ceratoma trifurcata*, freeze-tolerant pest), we also extracted information on the median survival time for individuals held at a constant temperature of 0°C (Lam and Pedigo 2000). Median survival time corresponds to the time required for 50% mortality in a sample population. To find this value, we fit a quadratic regression model to survival data published in Lam and Pedigo (2000). We then extracted the number of days corresponding to 50% mortality, as well as the associated confidence interval using the fitted regression line.

### Collection of Climate Data

To represent the range of variation in current and future winter conditions in the Great Lakes region, we installed active-warming greenhouses with automated, retractable roofs at nine sites throughout Minnesota, Wisconsin, and Michigan (Figure S1). Sites spanned a broad latitudinal gradient (42.9–46.8°N) and three habitats types: deciduous forest, coniferous forest, and open areas. At each site, we installed three greenhouses, each with a different temperature treatment: the control (GH_control_, internal temperature = ambient temperature), 3°C warmer than ambient (GH_+3°C_), and 5°C warmer than ambient (GH_+5°C_). We chose these treatments to mimic potential future climate scenarios, which predict warmer winter temperatures in the Great Lakes region with increases ranging from approximately 3.5–6°C (Notaro, et al. 2014). From December 2016–March 2017, we monitored ambient temperatures and wind speed at 1-minute intervals and subnivium temperature and snow depth at 5-minute intervals in each greenhouse. We also recorded the same factors outside of each greenhouse to understand current ambient conditions and to control for any effect of the greenhouse structure.

To capture subnivium changes in the different warming treatments, we paired each greenhouse with an external weather station that included a heated rain gauge and an ambient temperature sensor. Communication between the temperature instruments inside the greenhouse and this weather station allowed us to capture both precipitation events and future climate conditions through each greenhouse’s retractable roof and precise temperature control. For further details on the collection of climate data and the equipment used, see Thompson et al. 2021.

### Statistical Analysis

#### Subnivium Temperature

Each greenhouse was equipped with sixteen temperature sensors to measure the temperature in the subnivium (i.e., ground surface temperature). There were additionally twelve temperature sensors that measured subnivium temperatures external to the greenhouses. To derive a daily subnivium temperature for each treatment, we extracted the daily minimum ground temperature from each sensor for the period of December 1, 2016 to March 31, 2017 and then calculated the mean of those minimum temperatures for each treatment (environmental control, n = 12; greenhouse treatments, n = 16) and each day. We characterized subnivium conditions using daily minimum temperatures because these temperatures allow us to quantify the lower limit of what insects would have to endure during their overwintering.

#### Boosted Regression Trees

To generate regional predictions of ground temperatures, we followed the methodology of Thompson et al. (2021) and used boosted regression trees (BRT) to model daily minimum ground temperatures, with each treatment (external environment, GH_control_, GH_+3°C_, GH_+5°C_) modeled separately (Breiman 2001). Our predictors included daily maximum air temperature, daily minimum air temperature, daily median snow depth, daily mean snow density, daily mean wind speed, and habitat type. After identifying the optimal settings and number of iterations for the models (Supplementary Methods), we ran each model 50 times using the R package *dismo* (Hijmans, et al. 2011, R Core Team 2022). In each iteration, the data was split into training and testing sets (70% and 30% of the data, respectively), allowing us to account for both the stochasticity intrinsic to boosted regression tree models (Elith, et al. 2008) and the variability in our collected data. To assess the predictive performance of the models, we calculated predictive deviance and root mean square error for each iteration.

#### Spatial predictions

To predict subnivium temperatures across the broader Great Lakes region, we obtained spatially explicit data on air temperature, snow depth and density, wind speed, and land cover (Supplementary Methods). We predicted minimum ground temperatures across the Great Lakes region for each day of our study period (December 1, 2016–March 31, 2017) at a 1-km resolution for the environment external to the greenhouses, GH_control_, GH_+3°C_, and GH_+5°C_, with 50 bootstrap samples for each day/treatment combination. By comparing the predictions generated for the environment external to the greenhouses to those generated for GH_control_, we also identified an offset, or correction factor, so that the ground temperature predictions would be unbiased by any effects of the greenhouse structure (Supplementary Methods). We applied the correction factor to the predictions produced for GH_+3°C_ and GH_+5°C_, which left us with 50 predictive surfaces of minimum daily ground temperature for each of three scenarios: no winter warming, warming of 3°C, and warming of 5°C. Finally, we summarized the predictions for each scenario by finding the mean of the predicted minimum ground temperatures across the 50 predictive surfaces.

#### Quantifying Insect Vulnerability

To quantify the effect of future winter temperatures on the vulnerability of insects overwintering in the subnivium, we used the highest and lowest SCPs extracted from the literature for each selected species as a threshold and to represent the worst and best cases, respectively, for overwintering survival. Higher supercooling points indicate that a species is relatively more sensitive to cold temperatures since the species would experience either extracellular freezing (freeze-tolerance) or mortality (freeze-avoidant) before other species with lower supercooling points. We present the results for the highest SCP (i.e., worst case) here and direct the reader to the supplemental materials for results on the lowest SCP (i.e., best case). We applied each species-specific SCP as a threshold to the predictive surfaces generated for each climate scenario and assigned a 1 to cells with ground temperature values lower than the threshold (1 cell = 1 km^2^). Then, across each climate scenario and for each cell, we summed these instances to represent the number of days in the winter season that the minimum daily ground temperature fell below the species’ SCP. For the entire region, we also summed the cells that fell below each species’ threshold to represent the extent of vulnerability for each day in the winter season.

Although the number of days below the SCP provides a common metric with which to compare freeze-avoidant and freeze-tolerant species, the correlation between the number of days below the SCP and the vulnerability of freeze-tolerant species may not be as strong as in freeze-avoidant species, since freeze-tolerant species can survive at temperatures below their supercooling point (Bale and Worland 2005). Therefore, we also used data on the number of days until 50% percent mortality in bean leaf beetles (*C. trifurcata*, a freeze-tolerant pest) held at a constant temperature of 0°C. This mortality information provides an additional measure of vulnerability for this species, as well as a means of assessing the ability of the number of days below SCP to serve as a reliable indicator of vulnerability in a freeze-tolerant species. For the predictive surfaces generated for each climate scenario, we calculated the maximum consecutive days of daily minimum ground temperatures at or below 0°C in each grid cell. We then used the time until 50% mortality that was observed in a sample of *C. trifurcata* (± confidence interval) (Lam and Pedigo 2000) as a threshold to determine the spatial distribution of consecutive below-freezing days that were below, within, and above the range of consecutive days that *C. trifurcata* could withstand. Then, to compare the SCP-based vulnerability estimates with the mortality-based vulnerability estimates, we examined how the distribution of days with temperatures below the species’ SCP and the extent of vulnerability overlapped with these lower lethal ranges.

## Results

The boosted regression tree models performed well with predictive deviances ranging from 1.93–2.49 and RMSE values ranging from 1.39–1.58°C (Table S4, S5). Overall, we found that compared to current conditions, the total number of days below the SCP decreased for most species under warming of 3°C (Figure 1, S8, Table S6, S7). This was especially true in northwestern areas of the study region, where there were about 20 less days of sub-SCP conditions in the +3°C scenario, even though region-wide reductions ranged only from 1.4–6.3 days (Figure 1, 2, S9). Under warming of 5°C, the total number of days with temperatures below the species’ SCPs returned to those found under current conditions in central and southern areas, and surpassed those found under current conditions in northern regions, with species experiencing on average between 1.6 and 3.9 more days of sub-SCP conditions (Figure 2, S9, Table S6, S7). Notably, the low supercooling points of the two butterfly species (*P. canadensis* and *P.glaucus*, both freeze-avoidant pollinators) did not result in any days below the SCP in any of the warming scenarios (Figure 1, S8).

**Figure 1.**
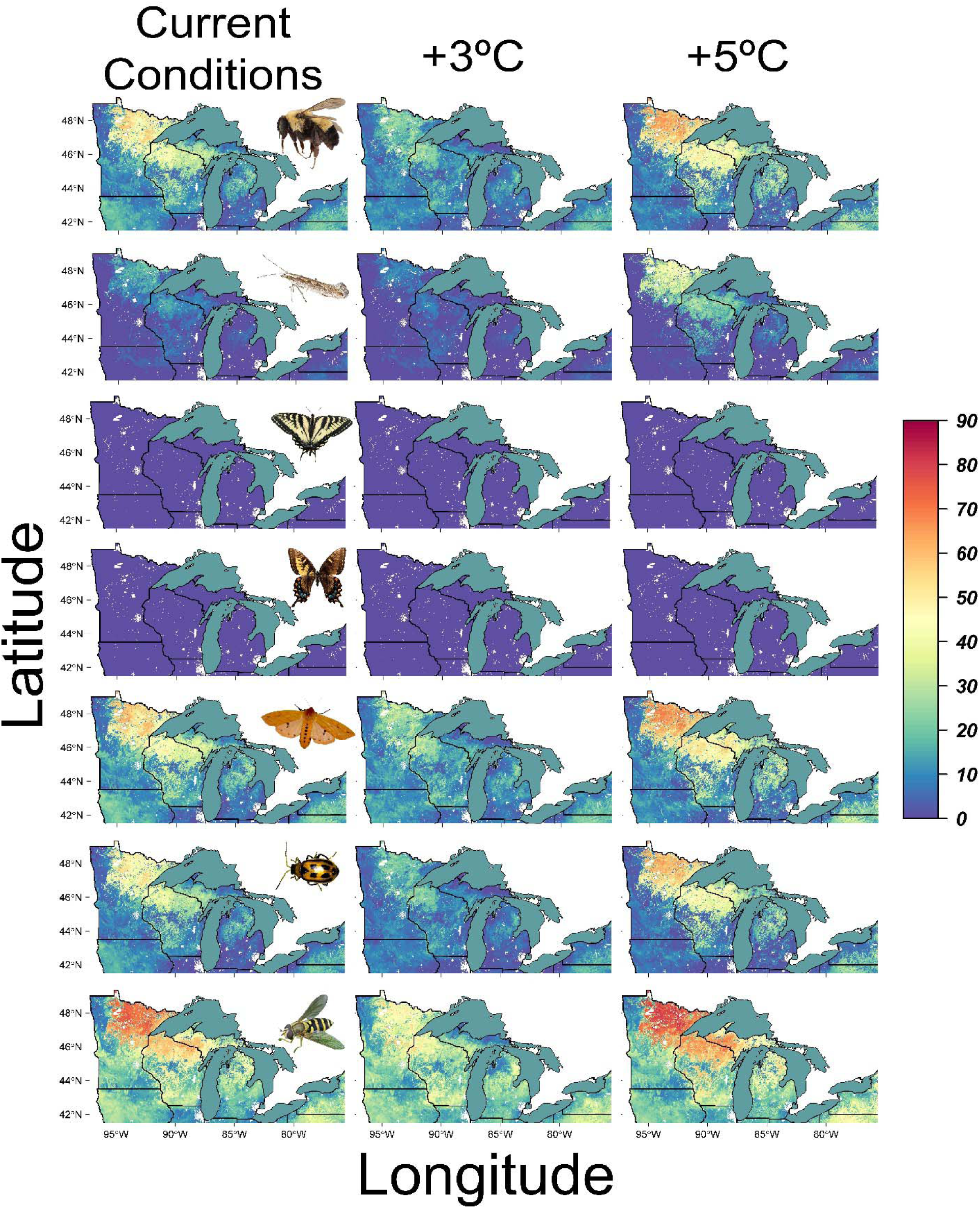
Total number of days in the winter season (December 1, 2016–March 31, 2017) below the highest published supercooling point (i.e., worst case) for insect species differing in their cold tolerance strategies and ecosystem services under current, 3°C warmer, and 5°C warmer conditions. From top to bottom: Buff-tailed bumblebee (*Bombus (Bombus) terrestris*, used a proxy for the Rusty patched bumblebee (*Bombus (Bombus) affinis*) and the yellow-banded bumblebee (*Bombus (Bombus) terricola*), freeze-avoidant pollinators; Diamondback moth (*Plutella xylostella*), freeze-avoidant pest; Canadian tiger swallowtail (*Papilio canadensis*), freeze-avoidant pollinator; Eastern tiger swallowtail (*Papilio glaucus*), freeze-avoidant pollinator; Woolly bear caterpillar (*Pyrrharctia isabella*), freeze-tolerant pollinator; Bean leaf beetle (*Ceratoma trifurcata*), freeze-tolerant pest; and Hoverfly (*Syrphus ribesii*), freeze-tolerant pollinator. Images of insects adapted from: *Rusty-patched bumblebee queen* by Miklasevskaja, M., 1971, https://val.vtecostudies.org/projects/vtbees/bombus-affinis/ Copyright 2024 by Vermont Center for Ecostudies; *Diamondback moth* 2006, https://en.wikipedia.org/wiki/Diamondback_moth; *Papilio canadensis* by Mdf, 2008, https://en.wikipedia.org/wiki/Papilio_canadensis; *Mosaic Gynandromorphs, Eastern Tiger Swallowtail (Papilio glaucus)* by Grace, K., 1979, https://www.floridamuseum.ufl.edu/100-years/object/eastern-tiger-swallowtail/ Copyright 2024 by Florida Museum of Natural History; *Pyrrharctia isabella* by Reago, A. and McClarren C., 2014 https://en.wikipedia.org/wiki/Pyrrharctia_isabella; *Adult bean leaf beetle* by University of Nebraska-Lincoln, 2024, https://cropwatch.unl.edu/soybean-management/insects-bean-leaf-beetle Copyright 1869-2024 by University of Nebraska-Lincoln; *Syrphus ribesii* by Aiwok, 2010, https://en.wikipedia.org/wiki/Syrphus_ribesii.

**Figure 2:**
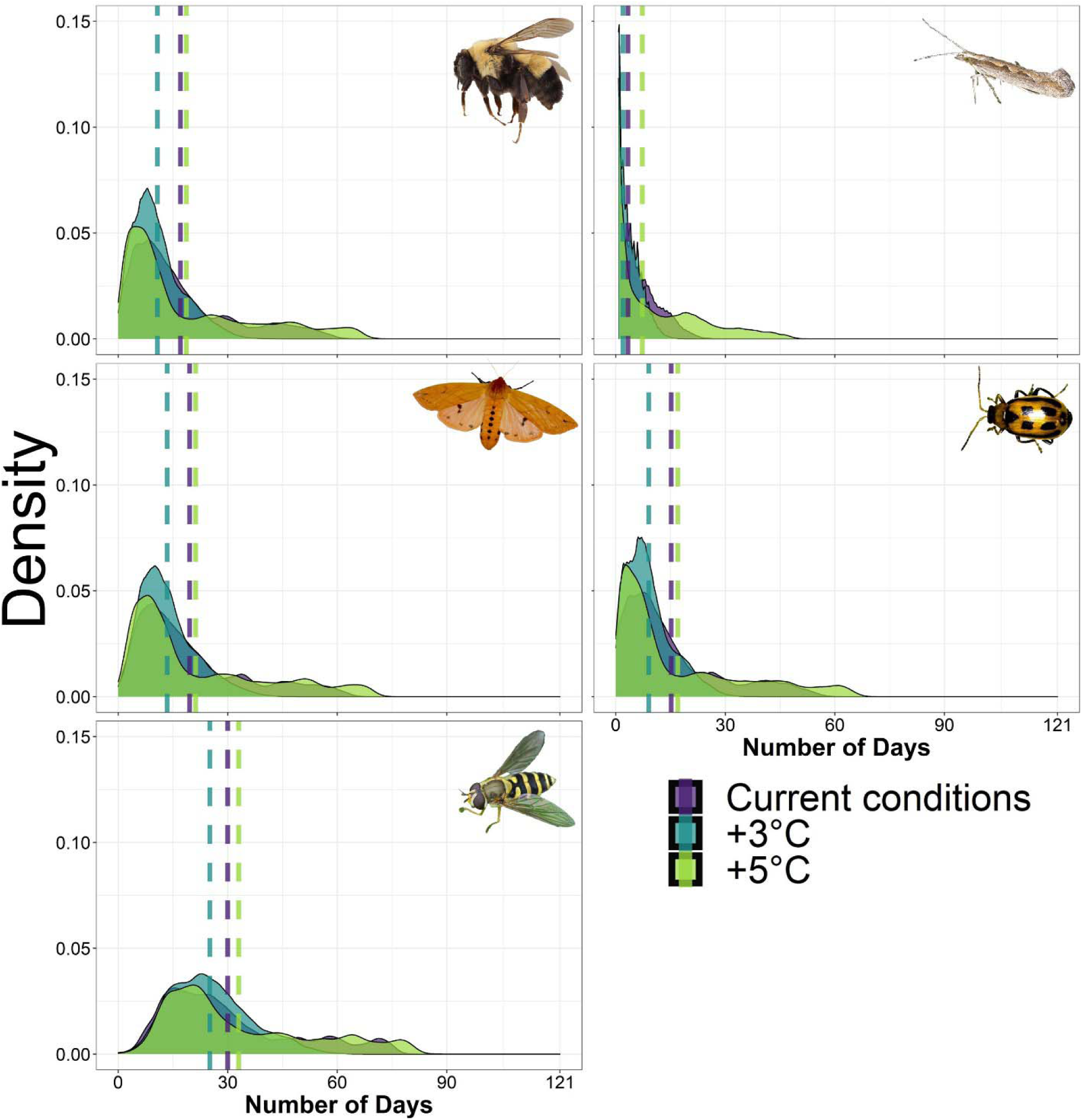
Density plots showing the distributions of the total number of days in the winter season (December 1, 2016–March 31, 2017) below the highest published supercooling point (i.e. worst case) for insect species differing in their cold tolerance strategies and ecosystem services under current, 3°C warmer, and 5°C warmer conditions. Dotted lines represent the mean number of days in each warming scenario. Top row: Buff-tailed bumblebee (*Bombus (Bombus) terrestris*, used a proxy for the Rusty patched bumblebee (*Bombus (Bombus) affinis*) and the yellow-banded bumblebee (*Bombus (Bombus) terricola*), freeze-avoidant pollinators, and the Diamondback moth (*Plutella xylostella*), freeze-avoidant pest. Middle row: Woolly bear caterpillar (*Pyrrharctia isabella*), freeze-tolerant pollinator and Bean leaf beetle (*Ceratoma trifurcata*), freeze-tolerant pest. Bottom row: Hoverfly (*Syrphus ribesii*), freeze-tolerant pollinator. The two butterfly species (Canadian tiger swallowtail (*Papilio canadensis*) and Eastern tiger swallowtail (*Papilio glaucus*, both freeze-avoidant pollinators) are not pictured because under each climate scenario the total days below each of their SCPs was zero. Images of insects adapted from: *Rusty-patched bumblebee queen* by Miklasevskaja, M., 1971, https://val.vtecostudies.org/projects/vtbees/bombus-affinis/ Copyright 2024 by Vermont Center for Ecostudies; *Diamondback moth* 2006, https://en.wikipedia.org/wiki/Diamondback_moth; *Pyrrharctia isabella* by Reago, A. and McClarren C., 2014 https://en.wikipedia.org/wiki/Pyrrharctia_isabella; *Adult bean leaf beetle* by University of Nebraska-Lincoln, 2024, https://cropwatch.unl.edu/soybean-management/insects-bean-leaf-beetle Copyright 1869-2024 by University of Nebraska-Lincoln; *Syrphus ribesii* by Aiwok, 2010, https://en.wikipedia.org/wiki/Syrphus_ribesii.

We found that the mean extent of vulnerability in the Great Lakes region improved under warming of 3°C, but expanded beyond the extents found under current conditions when warming reached 5°C (Table S8, S9). This increase in the extent of vulnerability between current conditions and warming of 5°C ranged from approximately 10,000 additional km^2^ for bumblebees (*B. affinus* and *B. terricola*, freeze-avoidant pollinators), woolly bear caterpillars *(P. isabella*, freeze-tolerant pollinator), and bean leaf beetles (*C. trifurcata*, freeze-tolerant pest) to 25,000 additional km^2^ for diamondback moths (*P. xylostella*, freeze-avoidant pest) and 80,000 additional km^2^ for hoverflies (*S. ribesii*, freeze-tolerant pollinator). Daily extents of vulnerability were highly variable throughout the winter season with many peaks and valleys (Figure 3, S10). In fact, despite the general pattern in which the mean extent of vulnerability increased under warming of 5°C, five species were predicted to experience a maximum daily extent of vulnerability under current conditions (*B. affinus*, *B. terricola*, *P. isabella*, *C. trifurcata*, and *S. ribesii*, Figure 3). The mean number of days until 50% mortality in a sample of bean leaf beetles held at a constant temperature of 0°C was 34.6 days [28.9, 41.3] (Figure S11; Lam and Pedigo 2000). Comparing this length of time to the number of consecutive days across the Great Lakes region with predicted daily minimum ground temperatures below 0°C, revealed additional areas of vulnerability for this species, as well as additional areas of improvement beyond those predicted by the SCP analysis (Figure 4). While current conditions throughout much of the western portion of the study area exceeded the number of below-freezing days *C. trifurcata* could withstand, the central and southern portions of these western states dramatically improved in the +3°C and +5°C scenarios. Despite local variation, the extent of vulnerability increased drastically across all warming scenarios when accounting for the consecutive number of below-freezing days, with an additional 310,000 km^2^ of vulnerability under current conditions, an additional135,000 km^2^ under warming of 3°C, and an additional 162,000 km^2^ under warming of 5°C (Figure 4, Table S10).

**Figure 3.**
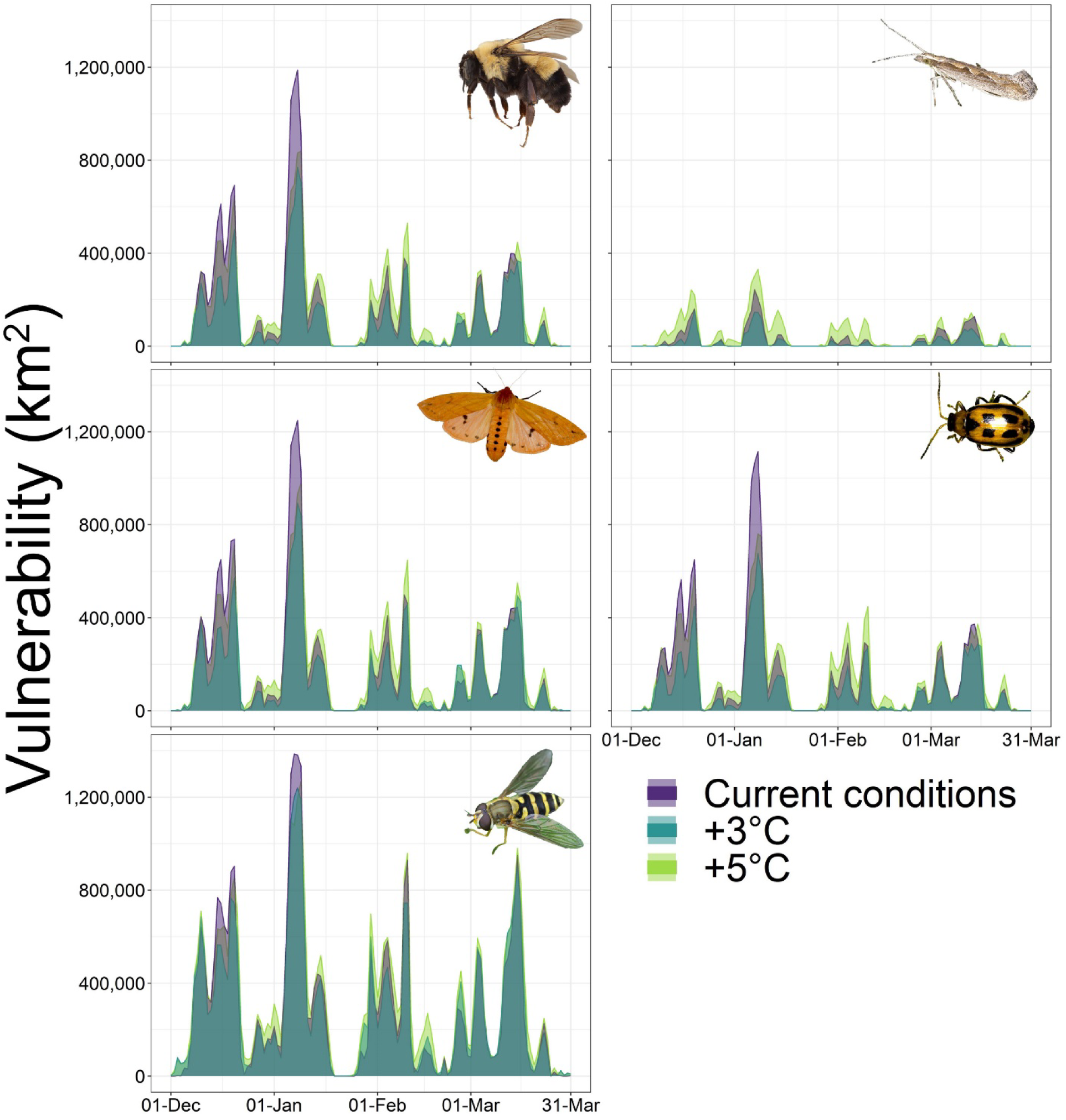
Daily extent of vulnerability (i.e., square kilometers where predicted ground temperatures were below the highest published supercooling point (i.e. worst case)) for insect species differing in their cold tolerance strategies and ecosystem services under current, 3°C warmer, and 5°C warmer conditions. Top row: Buff-tailed bumblebee (*Bombus (Bombus) terrestris*, used a proxy for the Rusty patched bumblebee (*Bombus (Bombus) affinis*) and the yellow-banded bumblebee (*Bombus (Bombus) terricola*), freeze-avoidant pollinators, and the Diamondback moth (*Plutella xylostella*), freeze-avoidant pest. Middle row: Woolly bear caterpillar (*Pyrrharctia isabella*), freeze-tolerant pollinator and Bean leaf beetle (*Ceratoma trifurcata*), freeze-tolerant pest. Bottom row: Hoverfly (*Syrphus ribesii*), freeze-tolerant pollinator. The two butterfly species (Canadian tiger swallowtail (*Papilio canadensis*) and Eastern tiger swallowtail (*Papilio glaucus*, both freeze-avoidant pollinators) are not pictured because under each climate scenario the total days below each of their SCPs was zero. Images of insects adapted from: *Rusty-patched bumblebee queen* by Miklasevskaja, M., 1971, https://val.vtecostudies.org/projects/vtbees/bombus-affinis/ Copyright 2024 by Vermont Center for Ecostudies; *Diamondback moth* 2006, https://en.wikipedia.org/wiki/Diamondback_moth; *Pyrrharctia isabella* by Reago, A. and McClarren C., 2014 https://en.wikipedia.org/wiki/Pyrrharctia_isabella; *Adult bean leaf beetle* by University of Nebraska-Lincoln, 2024, https://cropwatch.unl.edu/soybean-management/insects-bean-leaf-beetle Copyright 1869-2024 by University of Nebraska-Lincoln; *Syrphus ribesii* by Aiwok, 2010, https://en.wikipedia.org/wiki/Syrphus_ribesii.

**Figure 4.**
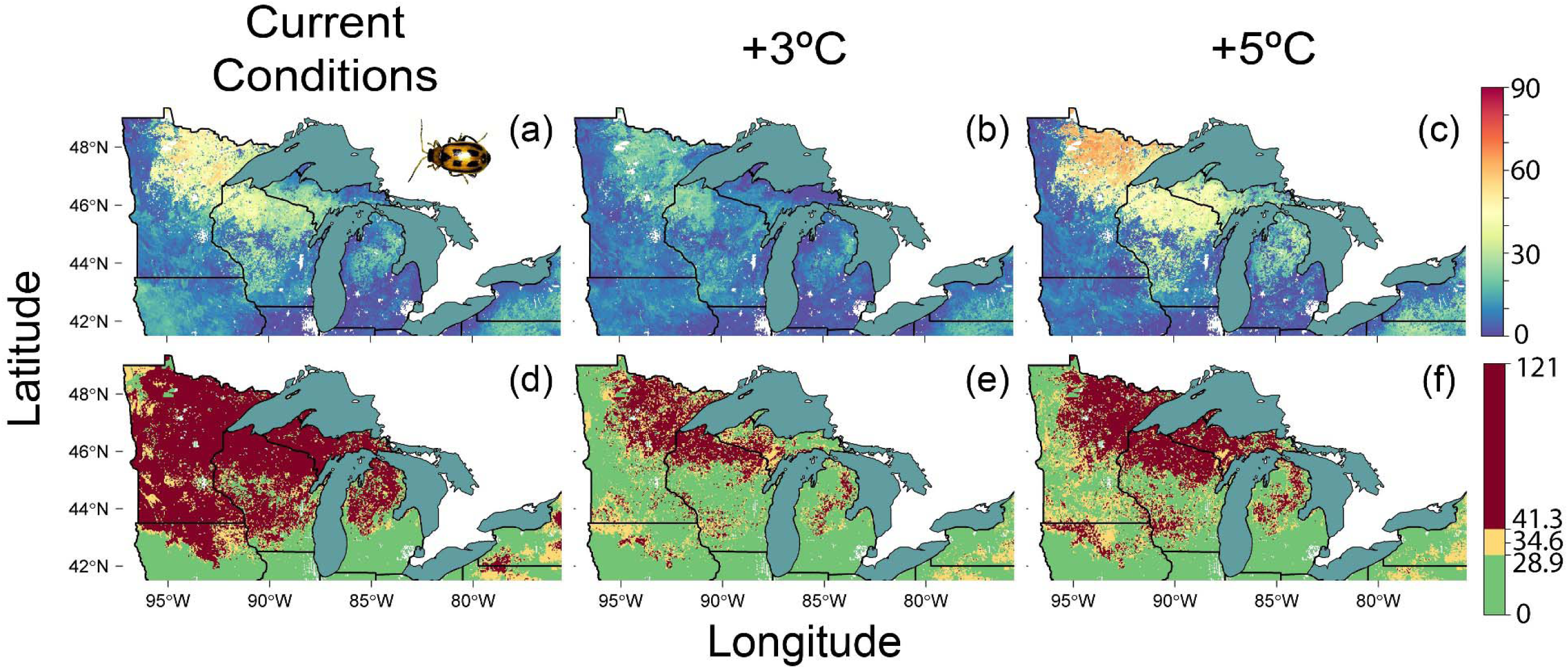
For three warming scenarios: current conditions (i.e. no warming, a & d), warming of 3°C (b & e), and warming of 5°C (c & f), comparison between the total number of days in the winter season (December 1, 2016–March 31, 2017) with ground temperatures below the highest published supercooling point (-7.0°C, i.e., worst case) for bean leaf beetles (*Ceratoma trifurcata*, freeze-tolerant pest; a–c) and mortality expectations based on experimental data extracted from Lam and Pedigo (2000) (d–f), who found that the mean number of days until 50% mortality in a sample of *C. trifurcata* at constant temperatures of 0°C was 34.6 days [28.9, 41.3]. For d–f, areas in red represent consecutive sub-0°C days exceeding the upper confidence limit of what *C. trifurcata* can withstand (i.e., least favorable scenario), areas in yellow represent consecutive sub-0°C days within the reported range for 50% mortality (i.e., moderate scenario), and areas in green represent sub-0°C days below the lower confidence limit of what *C. trifurcata* can withstand (i.e., most favorable scenario). Image of insect adapted from *Adult bean leaf beetle* by University of Nebraska-Lincoln, 2024, https://cropwatch.unl.edu/soybean-management/insects-bean-leaf-beetle Copyright 1869-2024 by University of Nebraska- Lincoln.

Comparing the distribution of total sub-SCP days with the lower lethal durations of sub- 0°C temperatures under current conditions and the +3°C climate scenario revealed considerable overlap between the three types of lower lethal durations (most favorable scenario: consecutive sub-0°C days below the duration that resulted in 50% mortality, moderate scenario: consecutive sub-0°C days within the duration that resulted in 50% mortality, and least favorable scenario: consecutive sub-0°C days above the duration that resulted in 50% mortality) when the total number of sub-SCP days ranged from 0–20 (Figure 5, S12). This indicates that mortality in bean leaf beetles is possible even when the number of days in which the temperature falls below the species SCP is low or zero. In the +5°C scenario, there was a marked increase in the number of sub-SCP days, leading to a concurrent increase in the number of consecutive sub-0°C days that were beyond what *C. trifurcata* could withstand (Figure 5, S12).

**Figure 5.**
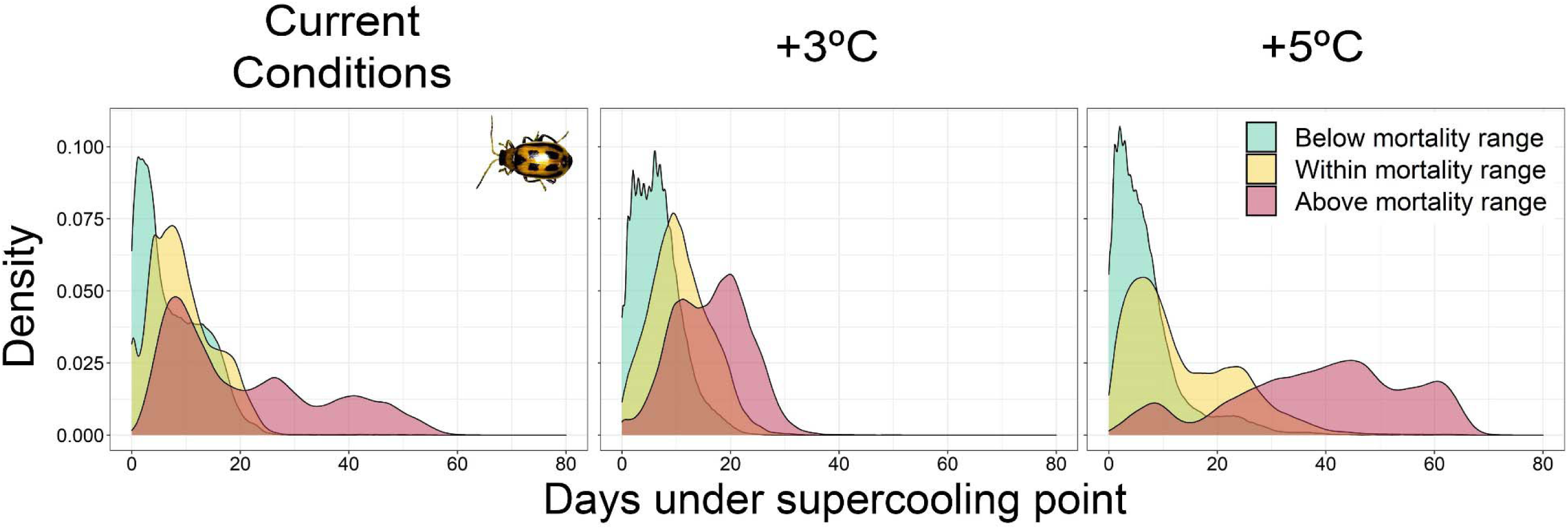
For three warming scenarios: current conditions (i.e. no warming), warming of 3°C, and warming of 5°C, density plots showing the overlap between the number of days during the winter season (December 1, 2016–March 31, 2017) with ground temperatures under the highest published supercooling point (-7.0°C, i.e., worst case) for bean leaf beetles (*C. trifurcata*) and the duration of consecutive days with ground temperatures at or below 0°C that would lead to 50% mortality based on experimental data extracted from Lam and Pedigo (2000). Lam and Pedigo (2000) found that the mean number of days until 50% mortality in a sample of *C. trifurcata* held at constant temperatures of 0°C was 34.6 days [28.9, 41.3]. ‘Above mortality range’ corresponds to consecutive sub-0°C days above this upper confidence limit of what *C. trifurcata* can withstand (i.e., least favorable scenario), ‘Within mortality range’ corresponds to consecutive sub-0°C days within the reported range for 50% mortality (i.e., moderate scenario), and ‘Below mortality range’ corresponds to sub-0°C days below the lower confidence limit of what *C. trifurcata* can withstand (i.e., most favorable scenario). Image of insect adapted from *Adult bean leaf beetle* by University of Nebraska-Lincoln, 2024, https://cropwatch.unl.edu/soybean-management/insects-bean-leaf-beetle Copyright 1869-2024 by University of Nebraska-Lincoln.

## Discussion

Warming winter temperatures are associated with a reduction in snow cover extent and a higher likelihood of rain-on-snow events, which in turn reduces the insulating capacity of snow cover (Zuckerberg and Pauli 2018). Consequently, even with warmer air temperatures, ground temperatures become paradoxically colder due to the lack of insulation (Brown and DeGaetano 2011). Despite this general phenomenon, prior work on the impact of winter warming on the spatial and temporal patterns of the subnivium demonstrated little to no change in extent or duration under warming of 3°C (Thompson, et al. 2021). Therefore, we hypothesized that species’ vulnerabilities would be roughly equivalent between the +3°C scenario and current conditions. Surprisingly, our results indicated that subnivium temperatures under warming of 3°C are likely to increase relative to current conditions, leading to reductions in the number of sub-SCP days and the mean extent of vulnerability for most insect species (Figure 1, 2, 3). Given that warming of 3°C does not generally involve deleterious effects for either the subnivium microhabitat or the insects that depend on it (Thompson, et al. 2021), efforts to reduce global emissions and limit warming could use this value as a critical benchmark for conserving the subnivium and communities of overwintering insects.

Under warming of 5°C, ground temperatures throughout the Great Lakes region decreased below those found in current conditions, which generally caused insect vulnerabilities to increase (Figure 1, 2, 3). There was considerable spatial variation in the magnitude of the change in vulnerability, however, with the northern areas of our study region becoming especially exposed to higher thermal extremes, while central and southern areas experienced more modest increases in vulnerability (Figure 1). Consequently, given both localized increases and decreases, the difference between the mean number of sub-SCP days for each species in the 5°C warming scenario and current conditions across the entire study region was low (Figure 2). Despite this geographic variation, the mean extent of variability between current conditions and warming of 5°C increased for all but the two butterfly species (*P. glaucus* and *P. canadensis*), indicating that warming of 5°C likely represents a tipping point at which exposure to lethal temperatures becomes much more difficult to avoid (Figure 3).

Notably, since prior work on the effects winter climate change in lake-effect areas predicted high probabilities of subnivium occurrence and long subnivium durations under warming of 5°C (Thompson, et al. 2021) due to reduced lake ice and more lake-effect snow (Notaro, et al. 2015), we expected low insect vulnerability in the northern areas surrounding the Great Lakes. Contrary to our hypotheses, these areas demonstrated some of the highest levels of vulnerability for insects under warming of 5°C (Figure 1, 4). Our results show that although continued lake-effect snow may partially safeguard subnivium extent and duration in these areas, warming of 5°C is likely to increase temperature variability within this microhabitat, thereby leading to increased vulnerability for overwintering insects.

Despite these general trends, species responses did not converge according to their cold tolerance strategies. For example, our predictions show that two freeze-avoidant butterfly species (*P. canadensis* and *P. glaucus)* will remain relatively buffered to changing subnivium conditions even under warming of 5°C due to their extremely low supercooling points (Figure 1). Similarly, the diamondback moth (*P. xylostella)*, a freeze-avoidant pest species, demonstrated extremely low sensitivity to both current conditions and warming of 3°C, although warming of 5°C was sufficient to increase its vulnerability in northern areas of our study region (Figure 1, 3). Alternatively, the situation was much bleaker for bumblebees (*Bombus spp.*) and hoverflies (*S. ribesii*) under both current conditions and warming of 5°C, despite the modest reductions in vulnerability under 3°C warming (Figure 1, 3).

Freeze-tolerant species have been regarded as more uniform in their responses to cold, since mortality occurs at temperatures below their supercooling point (Bale and Worland 2005). Critically, however, these species are not immune to an unlimited range of temperatures, nor are they necessarily able to survive extended periods of exposure to cold (Layne and Blakeley 2002). For example, although woolly bear caterpillars (*P. isabella*) can endure subzero temperatures, they are still susceptible to prolonged extreme cold outbreaks and demonstrate higher mortality during repeated freeze-thaw events (Layne, et al. 1999, Marshall and Sinclair 2011). Similarly, the freeze-tolerant bean leaf beetle (*C. trifurcata*) can only survive constant temperatures of 0°C for 29–41 days (Lam and Pedigo 2000), despite having a supercooling point around -8°C (Carrillo, et al. 2005). Our comparison of vulnerability in bean leaf beetles between this lower lethal limit (0°C for 29–41 days) and the number of days below its SCP (Figure 4, 5) confirmed this idea of a more cryptic vulnerability that is likely typical of other freeze-tolerant species since sustained below-freezing temperatures even at temperatures above the SCP were sufficient to induce vulnerability (i.e., predicted mortality). We found more areas of vulnerability for bean leaf beetles using the lower lethal limit than we did with the number of sub-SCP days, as well as overlap between the least favorable and moderate scenarios (i.e., consecutive sub-0°C days above 41 days, and within 29–41 days, respectively) and the number of winter season days below the bean leaf beetle’s SCP (worst-case SCP: -7.0°C, best-case SCP -9.3°C). Critically, this indicates that individuals of this species could experience mortality even in the absence of any days below their SCP (Figure 5, S12). These results echo those of Berzitis et al. (2017), who found extremely low overwintering survival rates of bean leaf beetles across experimental treatments in southern Canada that included warming of 4°C, complete snow removal with no warming, and intact snow cover with no warming, even though consecutive subfreezing days were uncommon in all but the snow removal treatment.

These nuances in interspecific vulnerability are especially important given the ecosystem services and disservices provided by insects, and the subsequent implications for both ecological processes and economic systems. Many insects provide critical pollination services, which contribute to the maintenance of genetic diversity in plant populations (Kearns, et al. 1998) and increase the yield of cultivated crops, with economic valuations in the hundreds of billions of dollars (Gallai, et al. 2009). Some pollinator species additionally help to control populations of pest species through predation (Hart and Bale 1998). Alternatively, pest species cause billions of dollars in damage to crops annually (Zalucki, et al. 2012), with biocontrol efforts like pesticide applications causing declines in other animals, including insect pollinators (Raine and Gill 2015). Given the similarities we found between pollinator and pest species, efforts to conserve pollinators will likely benefit pest species as well, highlighting the potential for ecosystem service and disservice trade-offs under future climate change.

Decreases in pollinator populations have been well-studied, with pesticides, parasites and pathogens, habitat fragmentation, and climate change among the most cited reasons for species declines (Cameron and Sadd 2020). Similarly, the drivers of insect pest outbreaks have also received considerable attention, and are most often attributed to land-use change (e.g., conversion to monoculture agriculture, urbanization) and climate change (Dale and Frank 2017). For both pollinators and pests, however, studies on the effects of climate change have focused primarily on the impacts of heat waves and drought (Brown, et al. 2016, Ju, et al. 2015), rather than the consequences of a deteriorating subnivium (but see for example Berzitis, et al. 2017, Huang 2017, Marshall and Sinclair 2012). Filling this gap in our knowledge is critical because overwintering survival and the fitness consequences incurred during overwintering likely represent important bottlenecks for the population dynamics of subnivium-dependent species (Woodard, et al. 2019).

### Caveats

Our experimental approach for the collection of climate data focused on utilizing active- warming greenhouses that were able to capture natural environmental variability in both winter precipitation (i.e., through the retractable roofs) and temperature (i.e., through communication between internal and external temperature sensors) (Thompson, et al. 2021). While this method offered advantages, our ability to capture climate variability was constrained by our observation period: the winter season of 2016–17. Although there is a high degree of uncertainty surrounding the expected frequency of extreme winter events like polar vortices in the future (Screen, et al. 2018), it is possible that warming of 3°C could lead to increased variability in temperature and precipitation beyond the scope of our experimental design (Schimanke, et al. 2013). Consequently, while our findings indicate a positive outlook for overwintering insects under warming of 3°C with reduced exposure to temperatures below their supercooling points, these results depend on the assumption that winter climate variability would resemble what was observed during the 2016–17 season.

Our predictions also do not account for phenotypic plasticity. Thermal tolerance can be plastic (Schou, et al. 2017); therefore, species with higher phenotypic plasticity in their thermal tolerance may be able to mitigate their responses to colder and more variable subnivium temperatures (Rodrigues and Beldade 2020). At the same time, evidence suggests that the magnitude of climate warming is likely to exceed the slightly broader tolerance ranges gained through phenotypic plasticity (Gunderson and Stillman 2015). Accordingly, phenotypic plasticity will not be wholly sufficient to buffer overwintering insects from changes in the subnivium.

## Conclusion

Cold tolerance is an essential component of insect fitness and one of the best determinants of species distributions (Andersen, et al. 2015); however, insect species exhibit considerable variation in their cold tolerance and overwintering strategies, as well as in their supercooling points and lower lethal temperatures (Sinclair, et al. 2003). Consequently, despite the general trends we found of reduced vulnerability under warming of 3°C and increased vulnerability under warming of 5°C, interspecific variation in response to shifting subnivium temperatures still exists. This variation in overwintering vulnerability under different climate change scenarios will impact conservation plans for pollinators and mitigation plans for pests. Uncovering these species-specific responses will require additional data on the lower range of insect thermal tolerances, since currently this information—especially for some critical pollinators (e.g. *Bombus spp.*)—is limited (Sinclair, et al. 2015). Equally important is the capacity to link thermal tolerance data with fine-scale data on microclimatic conditions. Through these efforts we can gain a better understanding of overwintering vulnerability for the diverse array of subnivium-dependent insects.

## Acknowledgements

We thank M. Moore for designing and developing the automated greenhouses and Operation Fresh Start for constructing them. We thank M. Fitzpatrick, S. Petty, A. Mosloff, and L. Werner II for their help in data collection, as well as B. Ace, J. Clare, J. Grauer, M. Garces-Restrepo, C. Latimer, C. Lane, and A. Shipley. We thank Dane County Parks, University of Wisconsin-Madison Arboretum, University of Wisconsin-Milwaukee Field Station, US Forest Service, University of Wisconsin-Steven’s Point, Hunt Hill Audubon Sanctuary, University of Minnesota Cloquet Forestry Center, and Michigan Technological University Ford Center for access to their land. We also thank C. Gratton, W.P. Porter, V. Radeloff and M. Turner for their feedback. Finally, we thank the University of Wisconsin Department of Forest and Wildlife Ecology for their support. Funding for the project was provided by National Science Foundation’s Macrosystems Biology program, grant EF-1340632.

## References

1. Andersen, J. L., et al. 2015. How to assess Drosophila cold tolerance: chill coma temperature and lower lethal temperature are the best predictors of cold distribution limits. - Functional Ecology 29: 55–65.

2. Andresen, J. A., et al. 2014. Historical Climate and Climate Trends in the Midwestern United States. - In: J. A. Winkler, et al. (eds), Climate Change in the Midwest: A Synthesis Report for the National Climate Assessment. Island Press.

3. Bale, J. and Worland, R. 2005. Insects and low temperature: from molecular biology to distributions and abundance. - Comparative Biochemistry and Physiology and Molecular & Integrative Physiology 141: S331–S331.

4. Beekman, M., et al. 1998. Diapause survival and post-diapause performance in bumblebee queens (Bombus terrestris). - Entomologia Experimentalis Et Applicata 89: 207–214.

5. Berzitis, E. A., et al. 2017. Winter warming effects on overwinter survival, energy use, and spring emergence of Cerotoma trifurcata (Coleoptera: Chrysomelidae). - Agricultural and Forest Entomology 19: 163–170.

6. Bokhorst, S., et al. 2012. Extreme winter warming events more negatively impact small rather than large soil fauna: shift in community composition explained by traits not taxa. - Global Change Biology 18: 1152–1162.

7. Breiman, L. 2001. Statistical modeling: The two cultures. - Statistical Science 16: 199–215.

8. Brown, C. L., et al. 2004. Freezing induces a loss of freeze tolerance in an overwintering insect. - Proceedings of the Royal Society B-Biological Sciences 271: 1507–1511.

9. Brown, M. J. F., et al. 2016. A horizon scan of future threats and opportunities for pollinators and pollination. - Peerj 4: e2249.

10. Brown, P. J. and DeGaetano, A. T. 2011. A paradox of cooling winter soil surface temperatures in a warming northeastern United States. - Agricultural and Forest Meteorology 151: 947–956.

11. Burnett, A. W., et al. 2003. Increasing Great Lake-effect snowfall during the twentieth century: A regional response to global warming? - Journal of Climate 16: 3535–3542.

12. Cameron, S. A., et al. 2007. A comprehensive phylogeny of the bumble bees (Bombus). - Biological Journal of the Linnean Society 91: 161–188.

13. Cameron, S. A. and Sadd, B. M. 2020. Global Trends in Bumble Bee Health. - In: A. E. Douglas (ed) Annual Review of Entomology, Vol 65. pp. 209–232.

14. Campbell, J. L., et al. 2009. Consequences of climate change for biogeochemical cycling in forests of northeastern North America. - Canadian Journal of Forest Research 39: 264–284.

15. Carrillo, M. A., et al. 2005. Supercooling point of bean leaf beetle (Coleoptera : Chrysomelidae) in Minnesota and a revised predictive model for survival at low temperatures. - Environmental Entomology 34: 1395–1401.

16. Changnon, S. A. and Jones, D. M. A. 1972. Review of influences of Great Lakes on weather. - Water Resources Research 8: 360–371.

17. Christopher, S. F., et al. 2008. The effect of soil freezing on N cycling: comparison of two headwater subcatchments with different vegetation and snowpack conditions in the northern Hokkaido Island of Japan. - Biogeochemistry 88: 15–30.

18. Dale, A. G. and Frank, S. D. 2017. Warming and drought combine to increase pest insect fitness on urban trees. - PloS one 12: e0173844.

19. Dancau, T., et al. 2018. Elusively overwintering: a review of diamondback moth (Lepidoptera: Plutellidae) cold tolerance and overwintering strategy. - Canadian Entomologist 150: 156–173.

20. Elith, J., et al. 2008. A working guide to boosted regression trees. - Journal of Animal Ecology 77: 802–813.

21. Fitzpatrick, M. F., et al. 2019. Modeling the distribution of niche space and risk for a freeze- tolerant ectotherm, *Lithobates sylvaticus*. - Ecosphere 10: e02788.

22. Gallai, N., et al. 2009. Economic valuation of the vulnerability of world agriculture confronted with pollinator decline. - Ecological Economics 68: 810–821.

23. Gao, Y., et al. 2015. Persistent cold air outbreaks over North America in a warming climate. - Environmental Research Letters 10: 044001.

24. Grillakis, M. G., et al. 2016. Climate-Induced Shifts in Global Soil Temperature Regimes. - Soil Science 181: 264–272.

25. Groffman, P. M., et al. 2001. Colder soils in a warmer world: A snow manipulation study in a northern hardwood forest ecosystem. - Biogeochemistry 56: 135–150.

26. Gunderson, A. R. and Stillman, J. H. 2015. Plasticity in thermal tolerance has limited potential to buffer ectotherms from global warming. - Proceedings of the Royal Society B: Biological Sciences 282: 20150401.

27. Hart, A. J. and Bale, J. S. 1998. Factors affecting the freeze tolerance of the hoverfly Syrphus ribesii (Diptera : Syrphidae). - Journal of Insect Physiology 44: 21–29.

28. Herrera, J. M., et al. 2018. Climatic niche breadth determines the response of bumblebees (Bombus spp.) to climate warming in mountain areas of the Northern Iberian Peninsula. - Journal of Insect Conservation 22: 771–779.

29. Hijmans, R. J., et al. 2011. Package ’dismo’. - In: Available online at: http://cran.r-project.org/web/packages/dismo/index.html.

30. Huang, J. 2017. Presence of snow coverage and its thickness affected the mortality of overwintering pupae of Helicoverpa armigera (Hubner) (Lepidoptera: Noctuidae). - International Journal of Biometeorology 61: 709–718.

31. Ju, R.-T., et al. 2015. Increases in both temperature means and extremes likely facilitate invasive herbivore outbreaks. - Scientific reports 5: 15715.

32. Kearney, M. and Porter, W. 2009. Mechanistic niche modelling: combining physiological and spatial data to predict species’ ranges. - Ecology Letters 12: 334–350.

33. Kearney, M. R. 2020. How will snow alter exposure of organisms to cold stress under climate warming? - Global Ecology and Biogeography 29: 1246–1256.

34. Kearns, C. A., et al. 1998. Endangered mutualisms: The conservation of plant-pollinator interactions. - Annual Review of Ecology and Systematics 29: 83–112.

35. Kelty, J. D. and Lee, R. E. 1999. Induction of rapid cold hardening by cooling at ecologically relevant rates in Drosophila melanogaster. - Journal of Insect Physiology 45: 719–726.

36. Kodra, E., et al. 2011. Persisting cold extremes under 21st-century warming scenarios. - Geophysical Research Letters 38: L08705.

37. Korslund, L. and Steen, H. 2006. Small rodent winter survival: snow conditions limit access to food resources. - Journal of Animal Ecology 75: 156–166.

38. Lam, W. K. F. and Pedigo, L. P. 2000. Cold tolerance of overwintering bean leaf beetles (Coleoptera : Chrysomelidae). - Environmental Entomology 29: 157–163.

39. Layne, J. R., et al. 1999. Cold hardiness of the woolly bear caterpillar (Pyrrharctia isabella Lepidoptera : Arctiidae). - American Midland Naturalist 141: 293–304.

40. Layne, J. R. and Blakeley, D. L. 2002. Effect of freeze temperature on ice formation and long- term survival of the woolly bear caterpillar (Pyrrharctia isabella). - Journal of Insect Physiology 48: 1133–1137.

41. Lee, R. E. 2010. A primer on insect cold-tolerance. - In: R. E. Lee and D. L. Denlinger (eds), Insect Low Temperature Biology. Cambridge University Press, pp. 3–34.

42. Lehikoinen, A., et al. 2011. The impact of climate and cyclic food abundance on the timing of breeding and brood size in four boreal owl species. - Oecologia 165: 349–355.

43. Lemke, P. and Ren, J. 2007. Observations: Changes in Snow, Ice and Frozen Ground.

44. Lenoir, J., et al. 2017. Climatic microrefugia under anthropogenic climate change: implications for species redistribution. - Ecography 40: 253–266.

45. Marchand, P. J. 2013. Life in the Cold: an introduction to winter ecology. - University Press of New England.

46. Marshall, K. E. and Sinclair, B. J. 2011. The sub-lethal effects of repeated freezing in the woolly bear caterpillar Pyrrharctia isabella. - Journal of Experimental Biology 214: 1205–1212.

47. Marshall, K. E. and Sinclair, B. J. 2012. Threshold temperatures mediate the impact of reduced snow cover on overwintering freeze-tolerant caterpillars. - Naturwissenschaften 99: 33–41.

48. Mercader, R. J. and Scriber, J. M. 2008. Asymmetrical thermal constraints on the parapatric species boundaries of two widespread generalist butterflies. - Ecological Entomology 33: 537–545.

49. Neven, L. G., et al. 1986. Overwintering adaptations of the stag beetle, *Ceruchus piceus*, removal of ice nucleators in the winter to promote supercooling. - Journal of Comparative Physiology B-Biochemical Systemic and Environmental Physiology 156: 707–716.

50. Notaro, M., et al. 2014. Twenty-First-Century Projections of Snowfall and Winter Severity across Central-Eastern North America. - Journal of Climate 27: 6526–6550.

51. Notaro, M., et al. 2015. Dynamically Downscaled Projections of Lake-Effect Snow in the Great Lakes Basin. - Journal of Climate 28: 1661–1684.

52. O’Connor, J. H. and Rittenhouse, T. A. G. 2016. Snow cover and late fall movement influence wood frog survival during an unusually cold winter. - Oecologia 181: 635–644.

53. Overgaard, J. and MacMillan, H. A. 2017. The Integrative Physiology of Insect Chill Tolerance. - In: D. Julius (ed) Annual Review of Physiology, Vol 79. pp. 187–208.

54. Pauli, J. N., et al. 2013. The subnivium: a deteriorating seasonal refugium. - Frontiers in Ecology and the Environment 11: 260–267.

55. Peacock, S. 2012. Projected Twenty-First-Century Changes in Temperature, Precipitation, and Snow Cover over North America in CCSM4. - Journal of Climate 25: 4405–4429.

56. PRISM Climate Group 2020. - Oregon State University, https://prism.oregonstate.edu data created 4 Feb 2014, accessed 29 Nov 2022:

57. Pruitt, W. O. 2005. Why and how to study a snowcover. - Canadian Field-Naturalist 119: 118–128.

58. R Core Team 2022. R: A language and environment for statistical computing. - R Foundation for Statistical Computing.

59. Raine, N. E. and Gill, R. J. 2015. Tasteless pesticides affect bees in the field. - Nature 521: 38–40.

60. Rodrigues, Y. K. and Beldade, P. 2020. Thermal plasticity in insects’ response to climate change and to multifactorial environments. - Frontiers in Ecology and Evolution 8: 271.

61. Schimanke, S., et al. 2013. Variability and trends of major stratospheric warmings in simulations under constant and increasing GHG concentrations. - Climate Dynamics 40: 1733–1747.

62. Schou, M. F., et al. 2017. Linear reaction norms of thermal limits in Drosophila: predictable plasticity in cold but not in heat tolerance. - Functional Ecology 31: 934–945.

63. Scott, R. W. and Huff, F. A. 1996. Impacts of the Great Lakes on regional climate conditions. - Journal of Great Lakes Research 22: 845–863.

64. Screen, J. A., et al. 2018. Polar Climate Change as Manifest in Atmospheric Circulation. - Current Climate Change Reports 4: 383–395.

65. Scriber, J. M., et al. 2012. Differential effects of short term winter thermal stress on diapausing tiger swallowtail butterflies (Papilio spp.). - Insect Science 19: 277–285.

66. Serreze, M. C. 2010. Understanding Recent Climate Change. - Conservation Biology 24: 10–17.

67. Sinclair, B. J., et al. 2003. Climatic variability and the evolution of insect freeze tolerance. - Biological Reviews 78: 181–195.

68. Sinclair, B. J., et al. 2015. An invitation to measure insect cold tolerance: Methods, approaches, and workflow. - J. Therm. Biol. 53: 180–197.

69. Szabo, T. I. and Pengelly, D. H. 1973. Over-wintering and emergence of *Bombus* (*Pyrobombus*) *impatiens* (Creson) (Hymenoptera-Apidae) in southern Ontario. - Insectes Sociaux 20: 125–132.

70. Thompson, K. L., et al. 2018. The phenology of the subnivium. - Environmental Research Letters 13: 064037.

71. Thompson, K. L., et al. 2021. The decline of a hidden and expansive microhabitat: the subnivium. - Frontiers in Ecology and the Environment fee.2337.

72. Toxopeus, J. and Sinclair, B. J. 2018. Mechanisms underlying insect freeze tolerance. - Biological Reviews 93: 1891–1914.

73. Ullrich, T. S. 2020. xyscan. - In: Available online at: https://rhig.physics.yale.edu/~ullrich/software/xyscan/.

74. Vavrus, S., et al. 2006. The behavior of extreme cold air outbreaks under greenhouse warming. - International Journal of Climatology 26: 1133–1147.

75. Woodard, S. H., et al. 2019. Diet and nutritional status during early adult life have immediate and persistent effects on queen bumble bees. - Conservation Physiology 7: coz048.

76. Zalucki, M. P., et al. 2012. Estimating the Economic Cost of One of the World’s Major Insect Pests, Plutella xylostella (Lepidoptera: Plutellidae): Just How Long Is a Piece of String? - Journal of Economic Entomology 105: 1115–1129.

77. Zuckerberg, B. and Pauli, J. N. 2018. Conserving and managing the subnivium. - Conservation Biology 32: 774–781.

