## Supplemental Methods, Tables, and Figures for "The vulnerability of overwintering insects to loss of the subnivium"

*Supplementary Methods*

### Insect Species

#### Bumblebees: *Bombus affinis* and *Bombus terricola*

With their large relative size and dense fur, bumblebees (*Bombus spp.*) are well adapted for activity in cool conditions (Heinrich 1993) and are found throughout temperate, alpine, and arctic zones in North America, Europe, and Asia (Goulson 2010). Typically in late summer/early fall, young bumblebee queens who have mated seek out hibernacula in loose, disturbed soil, burrow several centimeters, and hibernate for a period of 6-9 months (Alford 1975). The starting time of this hibernation varies between species, with some entering diapause as early as May in extremely cold climates and other remaining active during the winter in milder climates (Goulson 2010). While some insects have obligatory diapause periods (Moraiti and Papadopoulos 2017, Sgolastra, et al. 2010), the flexibility in *Bombus spp.* suggests a more facultative, freeze-avoidant strategy (Owen, et al. 2013). For those bumblebee species that do hibernate, emergence from diapause in the spring is governed by temperature and photoperiod (Alford 1969, Larrere, et al. 1993). Once queens emerge they find a suitable site at which to found a colony and then lay their first batch of eggs (Heinrich 1972). Therefore, bumblebee queens emerging from diapause in early spring are solely responsible for the success of new colonies. While studies that examine the cold tolerance of bumblebee queens are rare, Owen et al. (2013) included queens in their study on *Bombus (Bombus) terrestris* and found a mean supercooling point of -7.0 ± 0.3°C (Table 1). Although *B. terrestris* is distributed in Europe, it exists in the same subgenus as *B. affinis* and *B. terricola* (Cameron, et al. 2007), two species with distributional ranges in the Great Lakes Region (Grixti, et al. 2009). Therefore, we use the SCP data for *B. terrestris* as a proxy for the cold tolerance of *B. affinis* and *B. terricola*.

**Diamondback Moth: *Plutella xylostella***

Diamondback moths are a globally distributed pest species that cause billions of dollars in crop damage annually (Zhang, et al. 2015) and are resistant to insecticides (Furlong, et al. 2013). In temperate climates they produce between 4–6 generations (Harcourt 1954) and as many as 20 generations in equatorial climates (Dosdall, et al. 2011). Although there is no consensus about the capacity of diamondback moths to overwinter at more northerly latitudes and the life stage during which overwintering occurs (Dancau, et al. 2018, Idris and Grafius 1996, Park and Kim 2014), several studies have demonstrated overwintering success at northerly latitudes in larvae (Idris and Grafius 1996), pupae (Razumov 1970), and adults (Saito 1994), with individuals spending the winter on leaf litter or crop debris, often under snow (Gu 2009, Kimura and Fujimura 1988).

A similar lack of consensus exists regarding the specific cold tolerance strategy employed by diamondback moths, since reported supercooling points are highly variable and often are equivalent to or lower than the lower lethal temperature (Dancau, et al. 2018). However, since this species slows its development in response to decreasing temperature, becomes immediately active when temperatures return to favorable conditions, and continues to feed even at low temperatures (Kimura, et al. 1987, Liu, et al. 2002), it likely overwinters in a state of quiescence rather than diapause, and is either freeze avoidant or chill susceptible (Dancau, et al. 2018).

In 2018, Dancau et al. reviewed the existing knowledge on supercooling for diamondback moths in each of its life stages; however, methodologies and estimates were highly variable between studies. We therefore extracted SCP data from Park and Kim (2014), who measured

SCPs for pupae and adults with and without a rapid cold hardening treatment. Despite these acclimation differences, differences in measured SCPs between pupae were small, and nonexistent in adults (Table 1) (Park and Kim 2014).

**Tiger Swallowtail butterflies**: ***Papilio canadensis* and *Papilio glaucus***

The Canadian tiger swallowtail (*Papilio canadensis*) and the Eastern tiger swallowtail (*Papilio glaucus*) are sister pollinator species whose range boundaries overlap in a hybrid zone that stretches from central Minnesota, Wisconsin, and Michigan to New England (Scriber 2011). This hybrid zone represents the southern range limit of the more northerly distributed *P. canadensis* and the northern range limit of the more southerly distributed *P. glaucus*. Both species are freeze avoidant and overwinter in leaf litter at soil surface; however, *P. Canadensis* is univoltine with obligate diapause, whereas P. glaucus is bivoltine with facultative diapause (Hagen and Scriber 1989, Rockey, et al. 1987). The majority of previous work has shown that both species overwinter as pupa (Mercader and Scriber 2008, Williams, et al. 2012), although recently diapause induction for *P. glaucus* was demonstrated to be more flexible and capable of occurring in the larval stage as well (Ryan, et al. 2018).

Kukal et al. (1991) examined the cold tolerance in a population of *P. Canadensis* from Michigan and a population of *P. glaucus* from Georgia, both of which were exposed to winter conditions in situ in Michigan for approximately two months prior to supercooling measurements. Despite the geographic distance in the origin of these populations, there was little variability in the supercooling points between these two species (Table 1) (Kukal, et al. 1991).

**Isabella Tiger Moth: *Pyrrharctia isabella***

Isabella tiger moths (*Pyrrharctia isabella*) are found throughout North America and overwinter at the soil surface under leaf litter as final instar caterpillars (aka woolly bears) (Layne, et al. 1999). The species is freeze tolerant; in the fall they move to diapause locations and cease feeding (Marshall and Sinclair 2011). Since they do not feed again after spring emergence and pupation (Goettel and Philogene 1987), they rely solely on the energy reserves they have left after diapause. Consequently, their metabolic expenditures in the winter and subsequent survival are very tied to snow cover (Marshall and Sinclair 2011).

Marshall and Sinclair (2011) measured variability in SCPs throughout two different winter seasons for woolly bear caterpillars experiencing repeated freeze-thaw events. The SCPs of caterpillars remained extremely consistent between freezing events, demonstrating very low intra-annual variability; however, SCPs between years were highly variable (Table 1) (Marshall and Sinclair 2011). SCPs were also reported by Layne et al. (1999) who calculated them using 4–8 caterpillars collected at four different times during the fall season (Table 1).

**Bean leaf beetle: *Ceratoma trifurcata***

Bean leaf beetles are native to eastern North America and are a major pest of soybeans and other legumes, causing direct damage by feeding on plants and indirect damage through transmission of plant pathogens (Hunt, et al. 1995). The species is freeze tolerant and overwinters as adults in forest leaf litter or in crop residue, typically entering diapause in early fall and emerging in mid to late spring based on photoperiod and temperature cues (Berzitis, et al. 2017, Lam and Pedigo 2000, Loughran and Ragsdale 1986). The number of generations produced annually also varies according to temperature, with 2-3 generations per year in central and southern United States, and one per year in more northern regions (McCreary 2013, Pedigo 1994).

Carrillo et al. (2005) examined the cold tolerance of bean leaf beetles collected from a soybean field in Minnesota and found low variability in SCPs during winter months, with mean values ranging from -8°C to -8.9°C and 90% confidence intervals ranging from -6.4°C to -9.9°C (Table 1) (Carrillo, et al. 2005).

**Hoverflies: *Syrphus ribesii***

Hoverflies (*Syrphus ribesii*) are distributed throughout the northern hemisphere, and in North America range from Alaska to Mexico (Brown, et al. 2004). In addition to providing pollination services, they also help to control pest species of aphids by feeding on them prior to overwintering (Hart and Bale 1998). Hoverflies are freeze tolerant, and overwinter as third instar larvae in leaf litter (Brown, et al. 2004).

Hart and Bale (1998) examined the effects of different acclimation treatments on the supercooling point of hoverflies, and found that as the duration of acclimation increased, the supercooling point also increased, indicating a concurrent increase in ice-nucleating activity to aid in freeze tolerance (Table 1). Brown et al. (2004) found a similarly high supercooling point for this species (Table 1); however, they also demonstrated that after one freeze event, *S. ribesii* were able to depress their supercooling point as low as -28°C and switch to a freeze-avoidant strategy. Despite this evidence for a mixed cold tolerance strategy and an extremely low supercooling point after one freeze event, individuals experiencing more than one freeze event were less likely to survive, pupate, or emerge (Brown, et al. 2004).

**Identifying Optimal Settings for Boosted Regression Tree Models**

Using the training data from the environment external to the greenhouses, we tested different combinations of learning rates, tree complexities, and total number of trees to determine the optimal combination of these settings for our BRT models. We evaluated learning rates of 0.05, 0.01, 0.005, 0.001, and 0.0005; tree complexities of 1–7; and total number of trees ranging from 100 to 300,000 in 100-tree increments. The models generated from these combinations were then applied to the testing dataset to assess which combination led to the lowest predictive deviance (Figure S2). Since our tests of tree complexities of 1 and 2 produced models with less than 1000 trees, we excluded them from further consideration (Figure S2). Using the remaining candidate combinations of tree complexity and learning rate, we then tested different bag fraction values ranging from 0.25–0.85 at 0.05-increments with total trees ranging from 100–30000 at 100-tree increments (Figure S3).

**Comparing Optimal BRT Model Setting Combinations**

We tested how sensitive the data were to different model settings by calculating the predictive deviance and root mean square error (RMSE) across 1000 iterations of each setting combination. We compared the deviance values using a one-way ANOVA and Tukey HSD post hoc tests, but used Welch tests for the RMSE values, since the RMSE values did not have homogeneous variances. These tests demonstrated that as tree complexity increased both predictive deviance and RMSE decreased, with each successive increase in tree complexity resulting in significantly lower deviance and RMSE (Figures S4, S5, Table S1). To balance predictive ability with the desire to avoid overfitting, we used the settings associated with a tree complexity of 5 (learning rate: 0.005, bag fraction: 0.65). Although these settings were derived using only the data for the external environment, we used this combination of settings in the BRT models for all other treatments ($GH_{control}$, $GH_{+3°C}$, and $GH_{+5°C}$), since the conditions in the external environment captured a range of natural variation.

**Bootstrapping the BRT models**

Using the selected combination of settings (tree complexity: 5, learning rate: 0.005, bag fraction: 0.65), we next compared the predictive deviance and RMSE between different numbers of model iterations to explore minimizing computation time. We compared deviances and

RMSEs for 25, 50, 75, 100, and 150 iterations of the BRT model built with the data for the environment external to the greenhouses to those produced from 1000 iterations using a one-way ANOVA to test for differences in means and Kolmorgorov Smirnov tests to compare distributions between samples. For predictive deviance, none of the means were significantly different from the deviance of the 1000-iteration sample (Table S2, Figure S6). Similarly, when comparing the distribution of each sample, there were no significant differences (Table S3). For RMSE, while there were significant differences, the mean RMSE of both the 50-iteration sample and 100-iteration sample were not significantly different from that of the 1000-iteration sample (Figure S7). Consequently, for each treatment, we ran the BRT models with our selected optimal settings 50 times.

**Data Used in Spatial Predictions**

Daily minimum and maximum temperature data was retrieved from Daymet, a product that interpolates daily meteorological data from weather stations throughout the whole of North America (Thornton, et al. 2016). For snow depth and snow density, we used data from the National Operational Hydrologic Remote Sensing Center's Snow Data Assimilation system (SNODAS). SNODAS data are based on an energy and mass-balance model that uses observations from satellites, airborne platforms, and ground stations to provide spatially-explicit snow characteristics (Carroll, et al. 2001). Daily mean wind speed was acquired from the National Centers for Environmental Prediction's North American Regional Reanalysis (Mesinger, et al. 2006, National Hydrologic Remote Sensing Center 2004). All continuous predictors were obtained at a resolution 1 $km^2$, with the exception of wind speed which was at a resolution of ~32 km.

We obtained land cover data at a 1-km resolution from the National Land Cover Database and reclassified the data into deciduous, coniferous, open, urban, water, and unclassified cover classes (Homer, et al. 2012). To better represent the three habitat types we included in the BRT models, we aggregated deciduous and mixed forest and woody wetlands into the deciduous cover class, evergreen forest and scrub/shrub into the conifer cover class, and developed open space, barren land, grassland/herbaceous, pasture/hay, cultivated crops, and emergent herbaceous wetlands into the open class.

**Correcting for the Greenhouse Structure**

To control for any effect of the greenhouse structure on subnivium temperatures, we quantified a correction factor to apply to our predictions of daily minimum ground temperatures. We defined the correction factor as the difference between predicted subnivium temperatures in $GH_{control}$ and the environment external to the greenhouses since in the absence of the greenhouse structure these predictions should be identical:

$$CF = T_{GH_{control}} - T_{external}$$

where $CF$ is the correction factor and $T$ is the predicted daily minimum ground temperature. We then used this correction factor to offset the predicted probabilities for $GH_{+3°C}$ and $GH_{+5°C}$ accordingly:

$$T_{corrected_{GH_{+5°C}}} = T_{GH_{+5°C}} - CF$$

$$T_{corrected_{GH_{+3°C}}} = T_{GH_{+3°C}} - CF$$

where $T_{corrected}$ is the corrected daily minimum ground temperature. Thus, for each raster cell in each treatment and iteration, we were able to generate a predictive surface in which the effect of the greenhouse warming structure was negligible.

Table S1. Tukey HSD pairwise comparisons from a one-way ANOVA comparing the predictive deviance of different boosted regression tree model settings across 1000 iterations [$F(4, 4995) = 165.9$, $p < 0.001$] and pairwise comparisons from a Welch test comparing the root mean square error for different boosted regression tree model settings across 1000 iterations [$F(4, 2496.5) = 2442.4$, $p < 0.001$]. TC=tree complexity, LR=learning rate, and BF=bag fraction.

| Comparison Metric | Group 1 Settings | | | Group 2 Settings | | | p | Diff | Lower Bound | Upper Bound |
|---|---|---|---|---|---|---|---|---|---|---|
| Predictive Deviance | TC=4 | LR=0.01 | BF=0.60 | TC=3 | LR=0.01 | BF=0.55 | 0.000 | -0.150 | -0.186 | -0.114 |
| | TC=5 | LR=0.005 | BF=0.65 | TC=3 | LR=0.01 | BF=0.55 | 0.000 | -0.189 | -0.225 | -0.152 |
| | TC=6 | LR=0.01 | BF=0.80 | TC=3 | LR=0.01 | BF=0.55 | 0.000 | -0.265 | -0.302 | -0.229 |
| | TC=7 | LR=0.005 | BF=0.80 | TC=3 | LR=0.01 | BF=0.55 | 0.000 | -0.031 | -0.352 | -0.279 |
| | TC=5 | LR=0.005 | BF=0.65 | TC=4 | LR=0.01 | BF=0.60 | 0.031 | -0.039 | -0.075 | -0.002 |
| | TC=6 | LR=0.01 | BF=0.80 | TC=4 | LR=0.01 | BF=0.60 | 0.000 | -0.115 | -0.152 | -0.079 |
| | TC=7 | LR=0.005 | BF=0.80 | TC=4 | LR=0.01 | BF=0.60 | 0.000 | -0.166 | -0.202 | -0.129 |
| | TC=6 | LR=0.01 | BF=0.80 | TC=5 | LR=0.005 | BF=0.65 | 0.000 | -0.077 | -0.113 | -0.040 |
| | TC=7 | LR=0.005 | BF=0.80 | TC=5 | LR=0.005 | BF=0.65 | 0.000 | -0.127 | -0.164 | -0.091 |
| | TC=7 | LR=0.005 | BF=0.80 | TC=6 | LR=0.01 | BF=0.80 | 0.001 | -0.051 | -0.087 | -0.014 |
| Root Mean Square Error | TC=4 | LR=0.01 | BF=0.60 | TC=3 | LR=0.01 | BF=0.55 | 0.000 | --- | --- | --- |
| | TC=5 | LR=0.005 | BF=0.65 | TC=3 | LR=0.01 | BF=0.55 | 0.000 | --- | --- | --- |
| | TC=6 | LR=0.01 | BF=0.80 | TC=3 | LR=0.01 | BF=0.55 | 0.000 | --- | --- | --- |
| | TC=7 | LR=0.005 | BF=0.80 | TC=3 | LR=0.01 | BF=0.55 | 0.000 | --- | --- | --- |
| | TC=5 | LR=0.005 | BF=0.65 | TC=4 | LR=0.01 | BF=0.60 | 0.000 | --- | --- | --- |
| | TC=6 | LR=0.01 | BF=0.80 | TC=4 | LR=0.01 | BF=0.60 | 0.000 | --- | --- | --- |
| | TC=7 | LR=0.005 | BF=0.80 | TC=4 | LR=0.01 | BF=0.60 | 0.000 | --- | --- | --- |
| | TC=6 | LR=0.01 | BF=0.80 | TC=5 | LR=0.005 | BF=0.65 | 0.000 | --- | --- | --- |
| | TC=7 | LR=0.005 | BF=0.80 | TC=5 | LR=0.005 | BF=0.65 | 0.000 | --- | --- | --- |

| | | | | | |
|---|---|---|---|---|---|
| TC=7 LR=0.005 BF=0.80 | TC=6 LR=0.01 BF=0.80 | 0.000 | --- | --- | --- |

Table S2. Tukey HSD pairwise comparisons from a one-way ANOVA comparing the predictive deviance [$F(5, 1394) = 1.407$, $p = 0.22$] and a one-way ANOVA comparing root mean square error [$F(5, 1394) = 14.36$, $p < 0.001$] for different numbers of bootstrap samples for a boosted regression tree model. All bootstraps used model settings of 5 for tree complexity, 0.005 for learning rate, and 0.65 for bag fraction.

| Comparison Metric | Number of Bootstraps | Number of Bootstraps | p | Lower Bound | Upper Bound |
|---|---|---|---|---|---|
| Predictive Deviance | 1,000 | 100 | 0.704 | -0.044 | 0.134 |
| | 150 | 100 | 0.545 | -0.045 | 0.174 |
| | 25 | 100 | 0.995 | -0.155 | 0.225 |
| | 50 | 100 | 0.864 | -0.088 | 0.206 |
| | 75 | 100 | 0.992 | -0.156 | 0.103 |
| | 150 | 1,000 | 0.975 | -0.055 | 0.094 |
| | 25 | 1,000 | 1.000 | -0.182 | 0.162 |
| | 50 | 1,000 | 1.000 | -0.109 | 0.137 |
| | 75 | 1,000 | 0.343 | -0.173 | 0.030 |
| | 25 | 150 | 0.997 | -0.213 | 0.154 |
| | 50 | 150 | 1.000 | -0.144 | 0.133 |
| | 75 | 150 | 0.257 | -0.211 | 0.029 |
| | 50 | 25 | 0.999 | -0.184 | 0.232 |
| | 75 | 25 | 0.948 | -0.257 | 0.135 |
| | 75 | 50 | 0.619 | -0.240 | 0.070 |
| Root Mean Square Error | 1000 | 100 | 1.000 | -0.008 | 0.007 |
| | 150 | 100 | 0.199 | -0.002 | 0.017 |
| | 25 | 100 | 0.000 | -0.042 | -0.010 |
| | 50 | 100 | 0.771 | -0.007 | 0.018 |

| | | | | |
|---|---|---|---|---|
| 75 | 100 | 0.001 | -0.027 | -0.005 |
| 150 | 1000 | 0.007 | 0.001 | 0.014 |
| 25 | 1000 | 0.000 | -0.041 | -0.012 |
| 50 | 1000 | 0.561 | -0.004 | 0.016 |
| 75 | 1000 | 0.000 | -0.024 | -0.007 |
| 25 | 150 | 0.000 | -0.049 | -0.018 |
| 50 | 150 | 0.999 | -0.013 | 0.010 |
| 75 | 150 | 0.000 | -0.033 | -0.013 |
| 50 | 25 | 0.000 | 0.015 | 0.050 |
| 75 | 25 | 0.424 | -0.006 | 0.027 |
| 75 | 50 | 0.000 | -0.034 | -0.008 |

Table S3. Kolmorgov-Smirnov results comparing the distribution of predictive deviance and root mean square error values between different numbers of bootstrap samples for a boosted regression tree model. All bootstraps used model settings of 5 for tree complexity, 0.005 for learning rate, and 0.65 for bag fraction.

| Comparison Metric | Number of Bootstraps | Number of Bootstraps | D-statistic | p |
|---|---|---|---|---|
| Predictive Deviance | 25 | 1,000 | 0.118 | 0.886 |
| | 50 | 1,000 | 0.134 | 0.360 |
| | 75 | 1,000 | 0.130 | 0.189 |
| | 100 | 1,000 | 0.118 | 0.159 |
| | 150 | 1,000 | 0.104 | 0.117 |
| Root Mean Square Error | 25 | 1,000 | 0.453 | 0.000 |
| | 50 | 1,000 | 0.171 | 0.123 |
| | 75 | 1,000 | 0.311 | 0.000 |
| | 100 | 1,000 | 0.074 | 0.702 |
| | 150 | 1,000 | 0.204 | 0.000 |

Table S4. Predictive deviance values for the boosted regression tree models built for conditions external to the greenhouses (external conditions) and each greenhouse treatment (+0°C, +3°C, and +5°C) using 50 bootstraps per model. GH=greenhouse.

| BRT Model | Mean | Std. Dev. | Median | Minimum | Maximum |
|---|---|---|---|---|---|
| External conditions | 2.43 | 0.27 | 2.43 | 1.73 | 3.13 |
| $GH_{control}$ (+0°C) | 2.49 | 0.33 | 2.47 | 1.83 | 3.41 |
| $GH_{+3°C}$ | 1.93 | 0.24 | 1.93 | 1.47 | 2.55 |
| $GH_{+5°C}$ | 2.31 | 0.23 | 2.30 | 1.92 | 2.81 |

Table S5. Root mean square error values for the boosted regression tree models built for conditions external to the greenhouses (external conditions) and each greenhouse treatment (+0°C, +3°C, and +5°C) using 50 bootstraps per model. GH=greenhouse.

| BRT Model | Mean | Std. Dev. | Median | Minimum | Maximum |
|---|---|---|---|---|---|
| External conditions | 1.56 | 0.02 | 1.56 | 1.51 | 1.60 |
| $GH_{control}$ (+0°C) | 1.58 | 0.02 | 1.58 | 1.53 | 1.61 |
| $GH_{+3°C}$ | 1.39 | 0.03 | 1.39 | 1.30 | 1.43 |
| $GH_{+5°C}$ | 1.50 | 0.03 | 1.50 | 1.45 | 1.61 |

Table S6. Summary of the total number of days in the winter season (December 1, 2016–March 31, 2017) below the highest published supercooling point (i.e., worst case) for each climate scenario and for insect species differing in their cold tolerance strategies and ecosystem services.

| Species | Climate Scenario | Mean | Standard Deviation | Minimum | Maximum |
|---|---|---|---|---|---|
| *Bombus terrestris*[a] | current conditions | 17.1 | 13.9 | 0.0 | 76.0 |
| | +3°C | 10.8 | 6.9 | 0.0 | 74.0 |
| | +5°C | 18.7 | 17.9 | 0.0 | 82.0 |
| *Plutella xylostella* | current conditions | 3.3 | 4.9 | 0.0 | 49.0 |
| | +3°C | 1.9 | 2.9 | 0.0 | 28.0 |
| | +5°C | 7.2 | 11.1 | 0.0 | 57.6 |
| *Papilio canadensis* | current conditions | 0.0 | 0.0 | 0.0 | 0.0 |
| | +3°C | 0.0 | 0.0 | 0.0 | 0.0 |
| | +5°C | 0.0 | 0.0 | 0.0 | 0.0 |
| *Papilio glaucus* | current conditions | 0.0 | 0.0 | 0.0 | 0.0 |
| | +3°C | 0.0 | 0.0 | 0.0 | 0.0 |
| | +5°C | 0.0 | 0.0 | 0.0 | 0.0 |
| *Pyrrharctia isabella* | current conditions | 19.5 | 14.8 | 0.0 | 78.0 |
| | +3°C | 13.4 | 7.7 | 0.0 | 76.0 |
| | +5°C | 21.2 | 18.5 | 0.0 | 85.0 |
| *Ceratoma trifurcata* | current conditions | 15.2 | 13.2 | 0.0 | 72.0 |
| | +3°C | 9.0 | 6.3 | 0.0 | 71.0 |
| | +5°C | 17.0 | 17.3 | 0.0 | 80.0 |
| *Syrphus ribesii* | current conditions | 30.0 | 17.9 | 0.0 | 88.0 |
| | +3°C | 25.1 | 10.9 | 0.0 | 91.0 |
| | +5°C | 33.0 | 20.1 | 0.0 | 95.0 |

[a] Used as a proxy for *Bombus (Bombus) affinis* and *Bombus (Bombus) terricola*

Table S7. Summary of the total number of days in the winter season (December 1, 2016–March 31, 2017) below the lowest published supercooling point (i.e., best case) for each climate scenario and for insect species differing in their cold tolerance strategies and ecosystem services.

| Species | Climate Scenario | Mean | Standard Deviation | Minimum | Maximum |
|---|---|---|---|---|---|
| *Bombus terrestris*[a] | current conditions | 13.3 | 12.4 | 0.0 | 71.0 |
| | +3°C | 7.6 | 5.8 | 0.0 | 65.0 |
| | +5°C | 15.4 | 16.8 | 0.0 | 78.0 |
| *Plutella xylostella* | current conditions | 0.5 | 1.2 | 0.0 | 11.0 |
| | +3°C | 0.3 | 0.9 | 0.0 | 11.0 |
| | +5°C | 2.0 | 4.0 | 0.0 | 34.0 |
| *Papilio canadensis* | current conditions | 0.0 | 0.0 | 0.0 | 0.0 |
| | +3°C | 0.0 | 0.0 | 0.0 | 0.0 |
| | +5°C | 0.0 | 0.0 | 0.0 | 0.0 |
| *Papilio glaucus* | current conditions | 0.0 | 0.0 | 0.0 | 0.0 |
| | +3°C | 0.0 | 0.0 | 0.0 | 0.0 |
| | +5°C | 0.0 | 0.0 | 0.0 | 0.0 |
| *Pyrrharctia isabella* | current conditions | 1.4 | 2.6 | 0.0 | 30.0 |
| | +3°C | 1.0 | 1.9 | 0.0 | 22.0 |
| | +5°C | 4.7 | 8.2 | 0.0 | 50.0 |
| *Ceratoma trifurcata* | current conditions | 4.2 | 5.9 | 0.0 | 54.0 |
| | +3°C | 2.3 | 3.2 | 0.0 | 35.0 |
| | +5°C | 8.0 | 11.9 | 0.0 | 60.0 |
| *Syrphus ribesii* | current conditions | 9.2 | 10.2 | 0.0 | 65.6 |
| | +3°C | 5.1 | 4.8 | 0.0 | 58.5 |
| | +5°C | 12.3 | 15.2 | 0.0 | 73.0 |

[a] Used as a proxy for *Bombus (Bombus) affinis* and *Bombus (Bombus) terricola*

Table S8. Summary of the extent of vulnerability ($km^2$) across the Great Lakes Region in the winter season (December 1, 2016–March 31, 2017) for each climate scenario and for insect species differing in their cold tolerance strategies and ecosystem services based on the highest published supercooling point (i.e., worst case).

| Species | Climate Scenario | Mean | Standard Deviation | Minimum | Maximum |
|---|---|---|---|---|---|
| *Bombus terrestris*[a] | current conditions | 157,776 | 232,058 | 0 | 1,187,785 |
| | +3°C | 115,924 | 153,697 | 0 | 768,903 |
| | +5°C | 165,732 | 185,250 | 0 | 838,608 |
| *Plutella xylostella* | current conditions | 27,192 | 46,134 | 0 | 243,846 |
| | +3°C | 17,434 | 31,618 | 0 | 153,598 |
| | +5°C | 53,150 | 68,227 | 0 | 330,676 |
| *Papilio canadensis* | current conditions | 0 | 0 | 0 | 0 |
| | +3°C | 0 | 0 | 0 | 0 |
| | +5°C | 0 | 0 | 0 | 0 |
| *Papilio glaucus* | current conditions | 0 | 0 | 0 | 0 |
| | +3°C | 0 | 0 | 0 | 0 |
| | +5°C | 0 | 0 | 0 | 0 |
| *Pyrrharctia isabella* | current conditions | 181,938 | 253,630 | 0 | 1,249,016 |
| | +3°C | 147,126 | 185,729 | 0 | 892,528 |
| | +5°C | 193,702 | 210,996 | 0 | 977,819 |
| *Ceratoma trifurcata* | current conditions | 140,038 | 213,472 | 0 | 1,114,632 |
| | +3°C | 95,455 | 131,436 | 0 | 676,078 |
| | +5°C | 147,366 | 167,423 | 0 | 760,348 |
| *Syrphus ribesii* | current conditions | 238,764 | 320,290 | 0 | 1,386,489 |
| | +3°C | 271,631 | 283,732 | 0 | 1,239,694 |
| | +5°C | 320,793 | 295,027 | 0 | 1,267,464 |

[a] Used as a proxy for *Bombus (Bombus) affinis* and *Bombus (Bombus) terricola*

Table S9. Summary of the extent of vulnerability (km$^2$) across the Great Lakes Region in the winter season (December 1, 2016–March 31, 2017) for each climate scenario and for insect species differing in their cold tolerance strategies and ecosystem services based on the lowest published supercooling point (i.e., best case).

| Species | Climate Scenario | Mean | Standard Deviation | Minimum | Maximum |
|---|---|---|---|---|---|
| *Bombus terrestris*[a] | current conditions | 121,641 | 191,211 | 0 | 1,002,921 |
| | +3°C | 78,567 | 111,482 | 0 | 576,180 |
| | +5°C | 130,728 | 151,150 | 0 | 696,452 |
| *Plutella xylostella* | current conditions | 4,064 | 9,629 | 0 | 45,952 |
| | +3°C | 2,542 | 6,280 | 0 | 30,334 |
| | +5°C | 13,277 | 23,072 | 0 | 125,168 |
| *Papilio canadensis* | current conditions | 0 | 0 | 0 | 0 |
| | +3°C | 0 | 0 | 0 | 0 |
| | +5°C | 0 | 0 | 0 | 0 |
| *Papilio glaucus* | current conditions | 0 | 0 | 0 | 0 |
| | +3°C | 0 | 0 | 0 | 0 |
| | +5°C | 0 | 0 | 0 | 0 |
| *Pyrrharctia isabella* | current conditions | 11,829 | 22,748 | 0 | 103,590 |
| | +3°C | 8,459 | 17,301 | 0 | 93,304 |
| | +5°C | 32,731 | 45,105 | 0 | 211,405 |
| *Ceratoma trifurcata* | current conditions | 34,082 | 56,294 | 0 | 292,036 |
| | +3°C | 21,120 | 37,040 | 0 | 174,408 |
| | +5°C | 59,976 | 76,268 | 0 | 377,659 |
| *Syrphus ribesii* | current conditions | 81,549 | 132,789 | 0 | 717,495 |
| | +3°C | 49,811 | 75,486 | 0 | 389,301 |
| | +5°C | 98,672 | 119,007 | 0 | 573,848 |

[a] Used as a proxy for *Bombus (Bombus) affinis* and *Bombus (Bombus) terricola*

Table S10. Comparison of the extents of vulnerability (km$^2$) during the winter season (December 1, 2016–March 31, 2017) across three warming scenarios (current conditions, +3°C, and +5°C) for bean leaf beetles (*Ceratoma trifurcata*) based on the number of days under the species' highest published supercooling point (SCP) (i.e., worst case) and the duration of consecutive days at or below 0°C that led to 50% mortality. Lam and Pedigo (2000) showed that the mean number of days until 50% mortality at constant temperatures of 0°C was 34.6 days [28.9, 41.3]. 'Extent above mortality range' corresponds to consecutive sub-0°C days above this upper confidence limit, 'Extent within mortality range' corresponds to consecutive sub-0°C days within the reported range for 50% mortality, and 'Extent below mortality range' corresponds to sub-0°C days below the lower confidence limit.

| Climate Scenario | Mean Extent of vulnerability based on SCP | Extent below mortality range | Extent within mortality range | Extent above mortality range | Sum of 'within' and 'above' mortality range | Difference between extent of 'within' and 'above' mortality range, and mean extent of vulnerability based on SCP |
|---|---|---|---|---|---|---|
| current conditions | 140,038 | 218,473 | 80,678 | 369,847 | 450,525 | 310,487 |
| +3°C | 95,455 | 438,072 | 113,417 | 117,509 | 230,926 | 135,471 |
| +5°C | 147,366 | 359,197 | 125,094 | 184,707 | 309,801 | 162,435 |

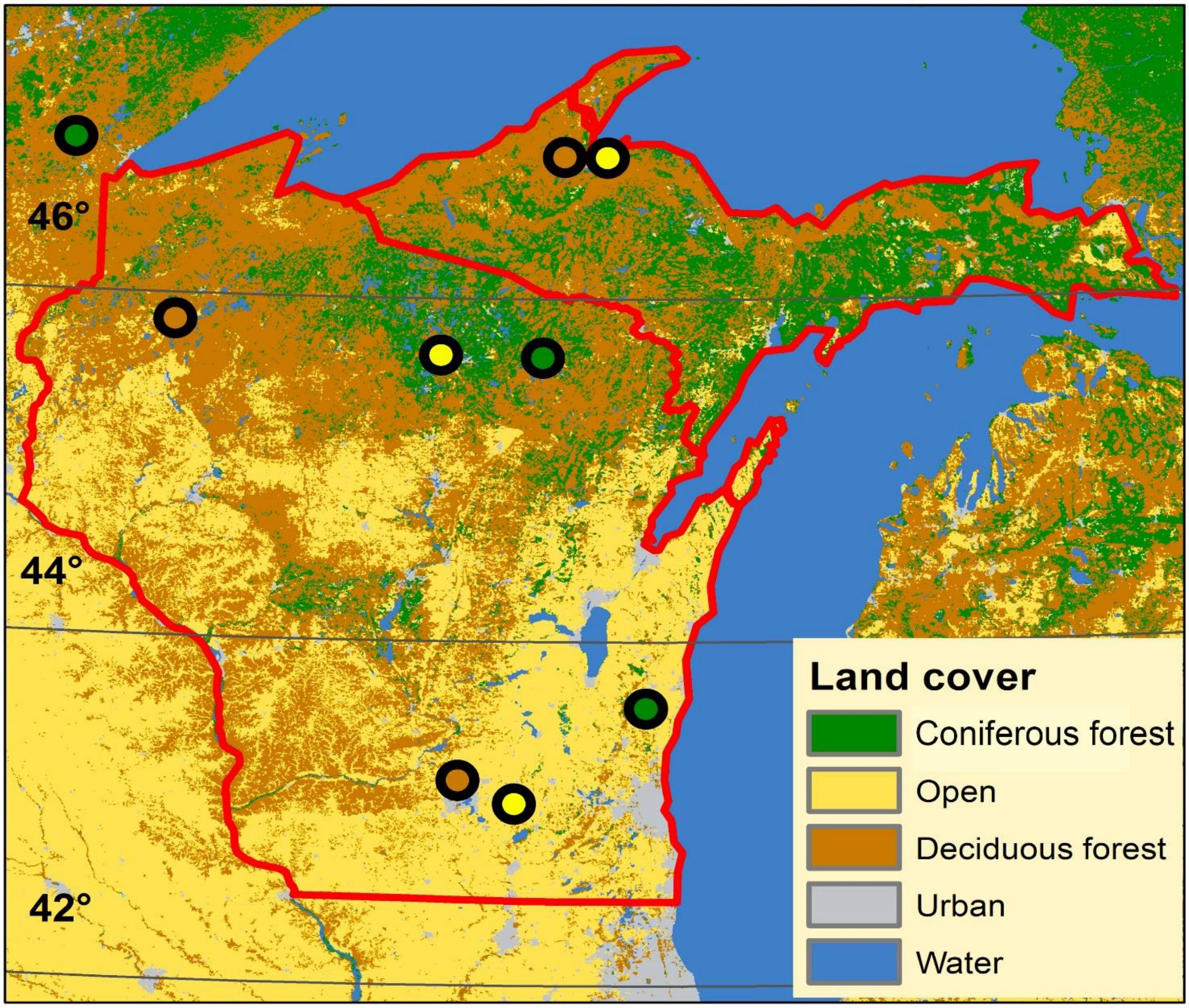


Figure S1. Active-warming greenhouses were installed at nine sites throughout the Great Lakes Region in three different habitat types (coniferous forest, deciduous forest, and open areas). Sites covered a broad latitudinal gradient to capture variation in winter temperatures and precipitation. At each site, we measured air temperature, subnivium temperature, snow depth, and wind speed in the external environment, a greenhouse with the same air temperature as the ambient temperature ($GH_{control}$), a greenhouse warmed to 3°C above ambient temperature ($GH_{+3°C}$), and a greenhouse warmed to 5°C above ambient temperature ($GH_{+5°C}$).

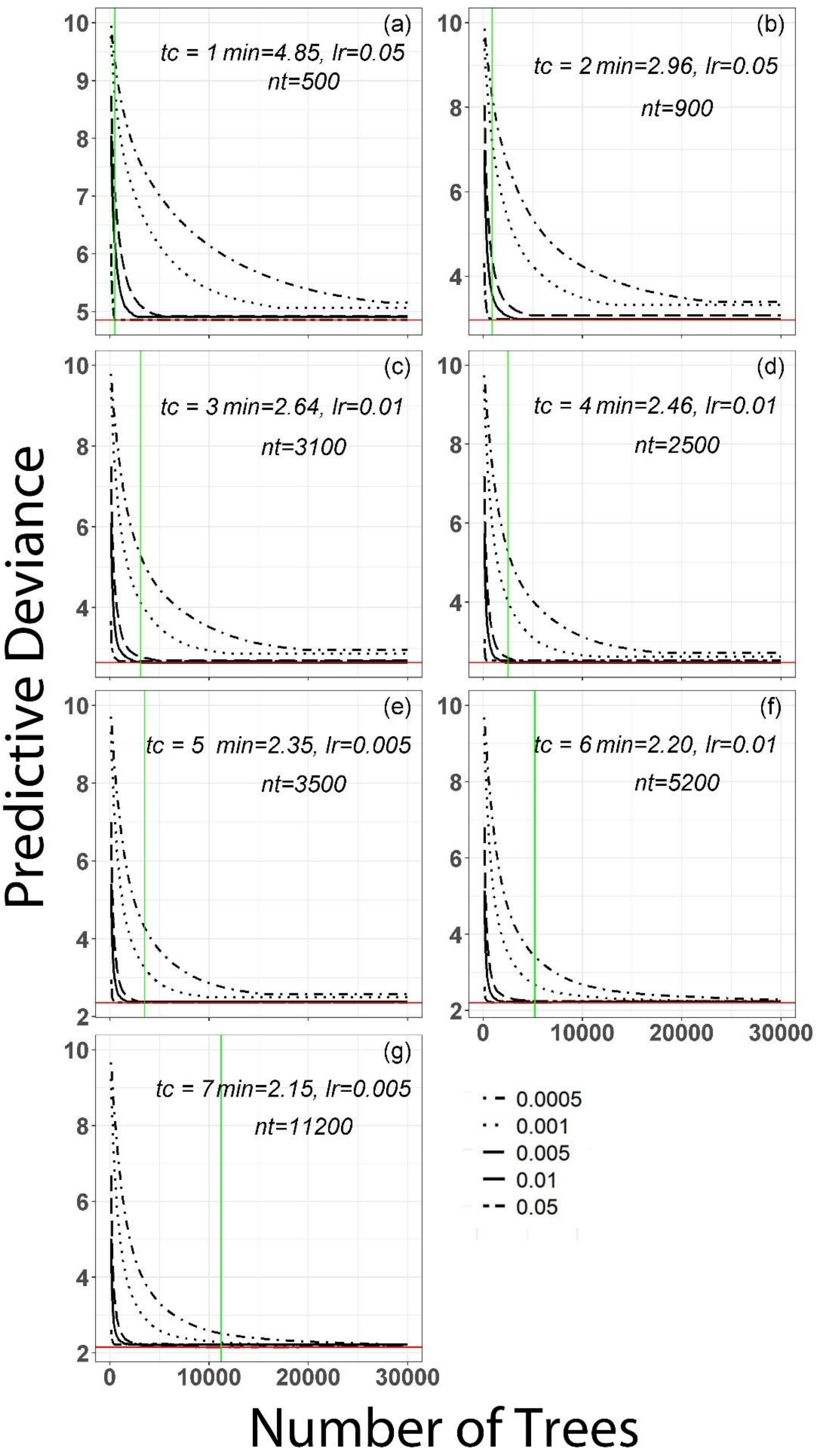
Predictive Deviance
Number of Trees
(a)
tc = 1 min=4.85, lr=0.05
nt=500
(b)
tc = 2 min=2.96, lr=0.05
nt=900
(c)
tc = 3 min=2.64, lr=0.01
nt=3100
(d)
tc = 4 min=2.46, lr=0.01
nt=2500
(e)
tc = 5 min=2.35, lr=0.005
nt=3500
(f)
tc = 6 min=2.20, lr=0.01
nt=5200
(g)
tc = 7 min=2.15, lr=0.005
nt=11200
0.0005
0.001
0.005
0.01
0.05
0
10000
20000
30000

244

Figure S2. Predictive deviance associated with different combinations of learning rate and number of trees for tree complexities of 1 through 7 (a through g, respectively). We tested learning rates of 0.05, 0.01, 0.005, 0.001, and 0.0005 (shown in the legend). The green line in each plot corresponds to the number of trees with which the lowest predictive deviance was found. The value of the lowest deviance is shown in each panel (min) along with the settings that produced this value: the tree complexity (tc), learning rate (lr), and number of trees (nt).

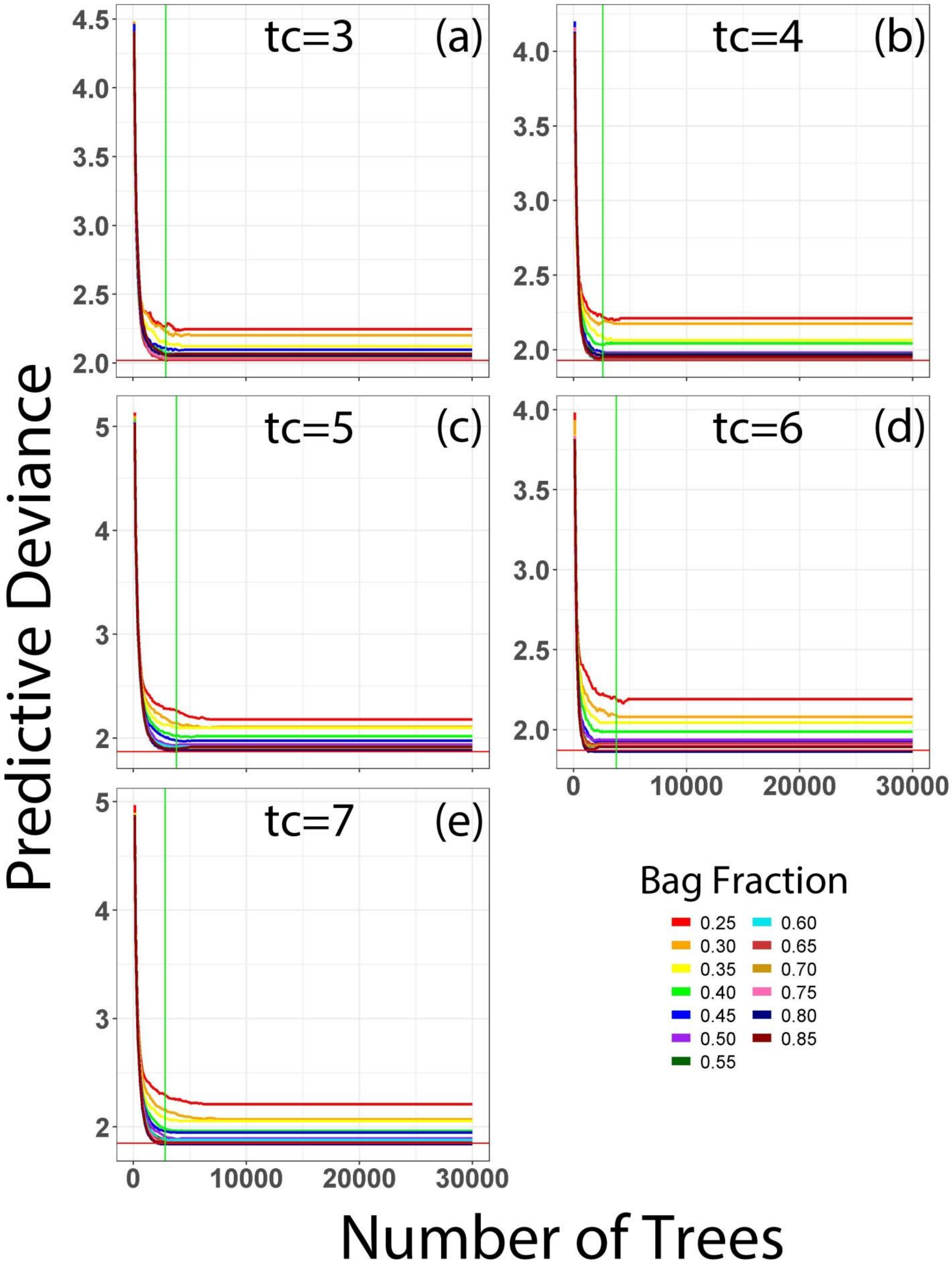

tc=3
(a)
tc=4
(b)
tc=5
(c)
tc=6
(d)
tc=7
(e)
Predictive Deviance
Number of Trees
Bag Fraction
0.25
0.30
0.35
0.40
0.45
0.50
0.55
0.60
0.65
0.70
0.75
0.80
0.85


268

Figure S3. Using tree complexities of 3–7 (a–e, respectively) and their corresponding learning rates (see Figure S1), we tested bag fractions (i.e. stochasticity values) ranging from 0.25–0.85 in increments of 0.05. We applied the models built from these different bag fractions to our testing dataset for the environment external to the greenhouses to determine the optimal bag fraction value for each tree complexity, based on which bag fraction resulted the lowest predictive deviance (represented by the intersection between the green vertical line and the red horizontal line).

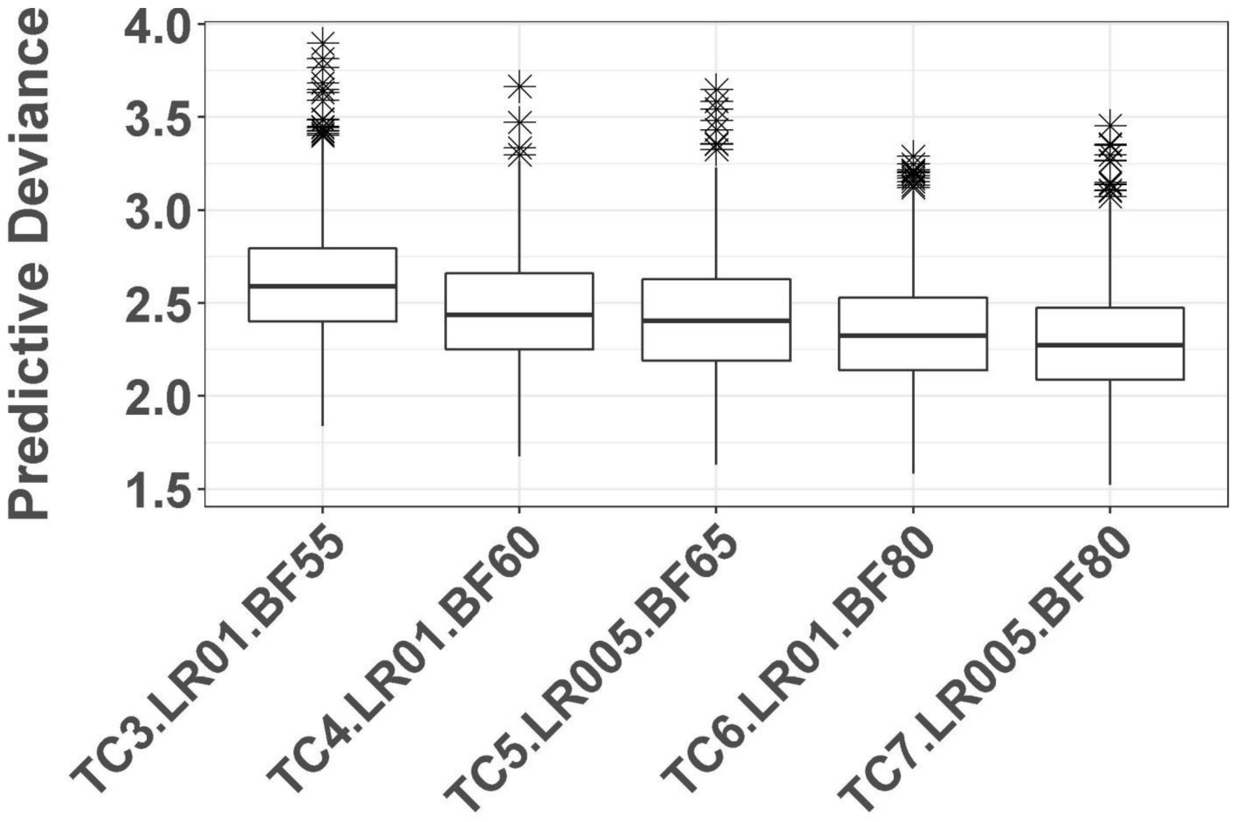


Figure S4. We tested how sensitive our data were to different boosted regression tree model settings by calculating the predictive deviance across 1000 iterations of each settings combination. TC=tree complexity and ranges from 3–7, LR=learning rate and is either 0.01 or 0.005, and BF=bag fraction and is either 0.55, 0.60, 0.65, or 0.80. As tree complexity increases predictive deviance decreases, with each successive increase in tree complexity resulting in significantly lower deviance.

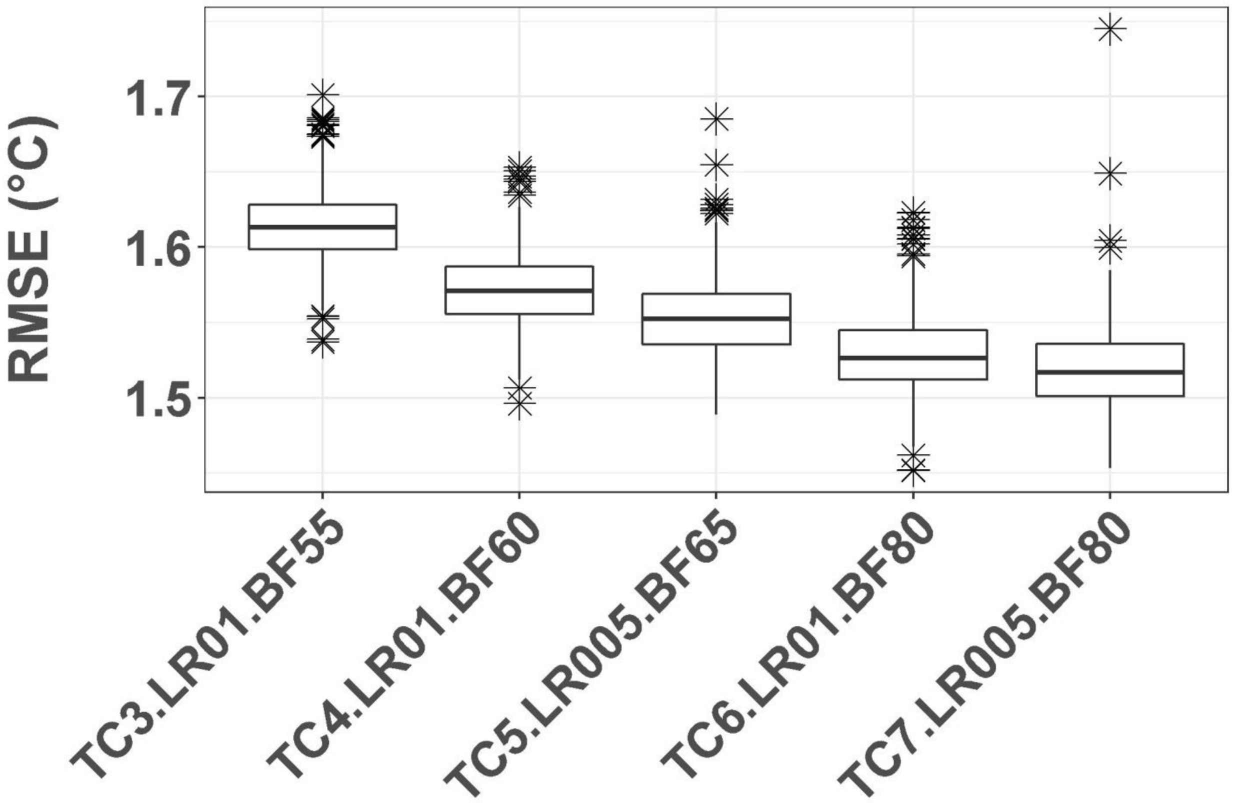


Figure S5. We tested how sensitive our data were to different boosted regression tree model settings by calculating the root mean square error (RMSE) across 1000 iterations of each settings combination. TC=tree complexity and ranges from 3–7, LR=learning rate and is either 0.01 or 0.005, and BF=bag fraction and is 0.55, 0.60, 0.65, or 0.80. As tree complexity increases RMSE decreases, with each successive increase in tree complexity resulting in significantly lower RMSE.

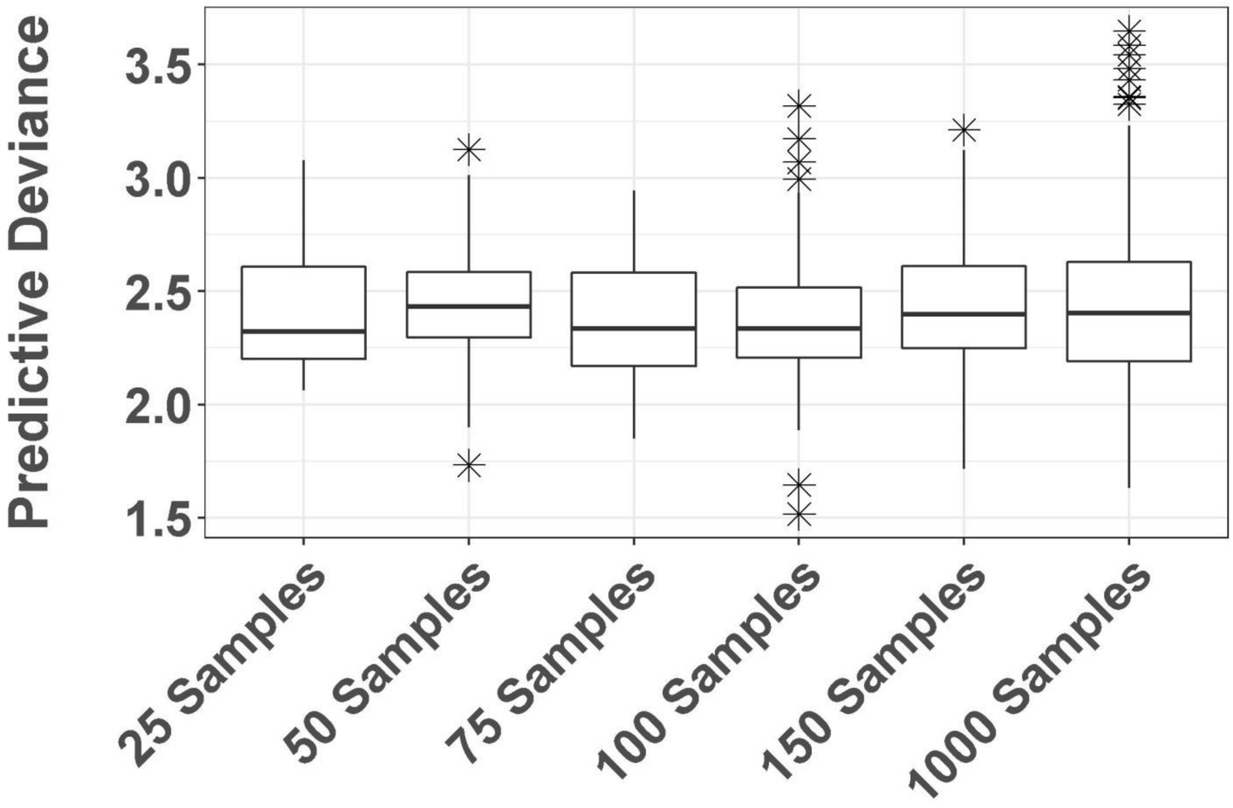


Figure S6. We tested how sensitive our data were to different numbers of bootstrap samples by calculating the predictive deviance across 25, 50, 75, 100, 150, and 1000 iterations of the boosted regression tree model. All bootstrap samples used model settings of 5 for tree complexity, 0.005 for learning rate, and 0.65 for bag fraction. None of the samples were significantly different from the deviance of the 1000-iteration sample.

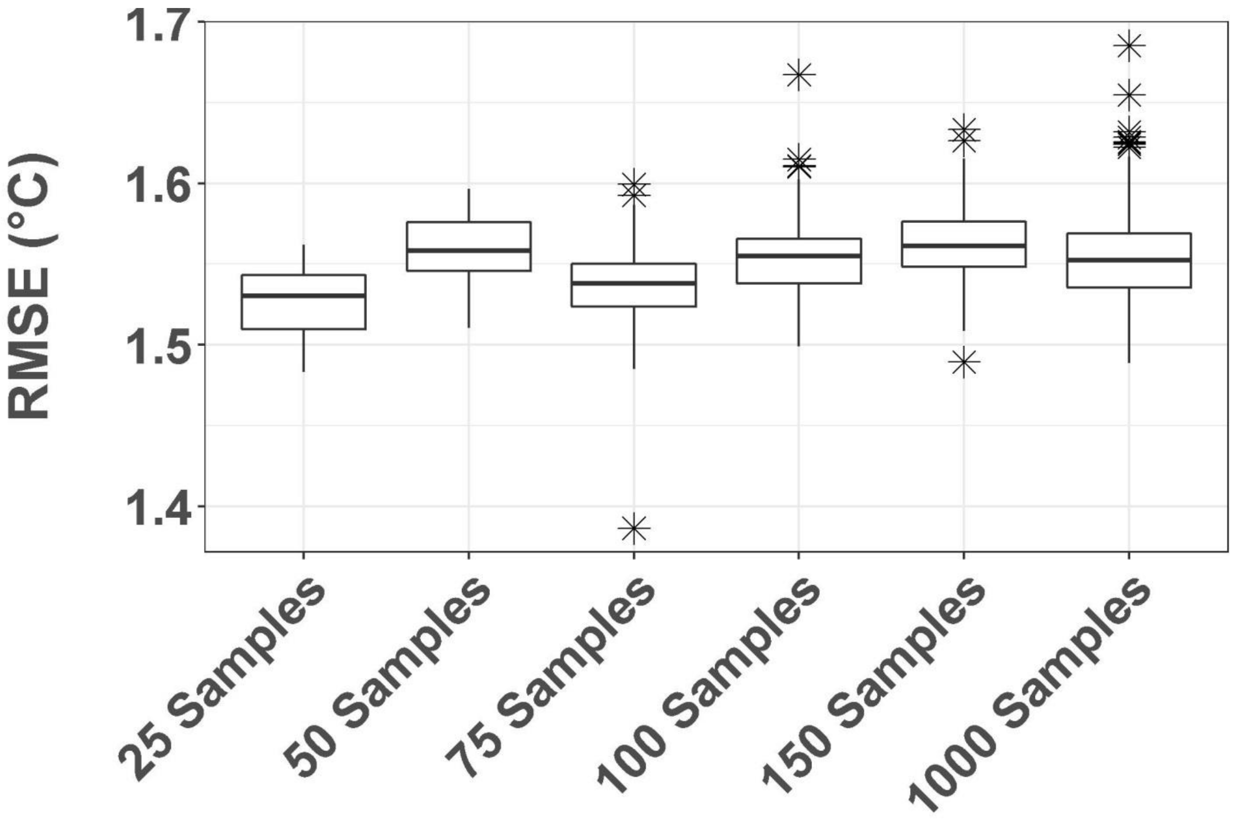


Figure S7. We tested how sensitive our data were to different numbers of bootstrap samples by calculating the root mean square error (RMSE) across 25, 50, 75, 100, 150, and 1000 iterations of the boosted regression tree model. All bootstrap samples used model settings of 5 for tree complexity, 0.005 for learning rate, and 0.65 for bag fraction. While there were significant differences, the mean RMSE of both the 50-bootstrap sample and 100-bootstrap sample were not significantly different from that of the 1000-bootstrap sample.

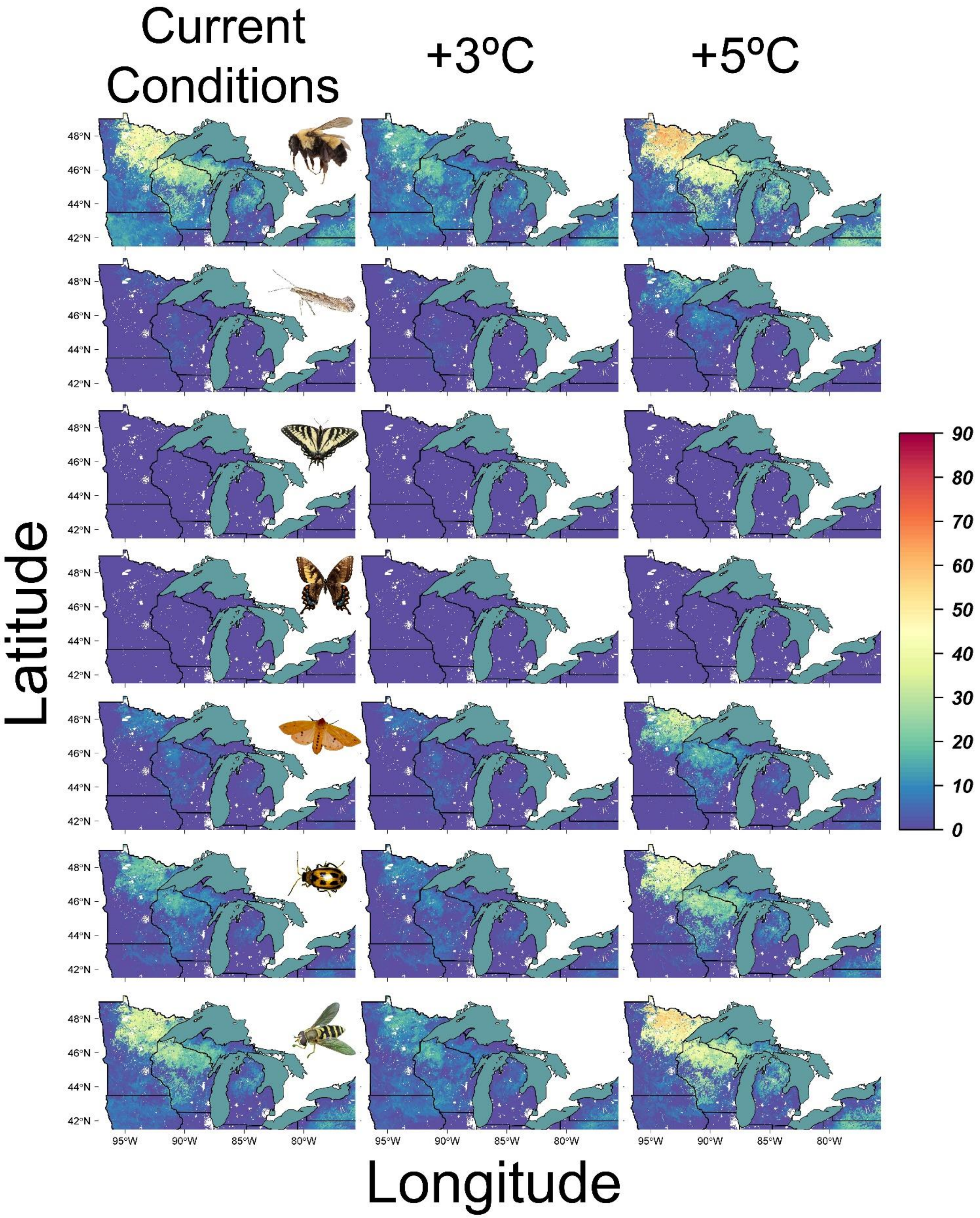

Current Conditions
+3ºC
+5ºC
Latitude
Longitude
48°N
46°N
44°N
42°N
95°W
90°W
85°W
80°W
90
80
70
60
50
40
30
20
10
0


332

Figure S8. Total number of days in the winter season (December 1, 2016–March 31, 2017) below the lowest published supercooling point (i.e., best-case scenario) for insect species differing in their cold tolerance strategies and ecosystem services under current, 3°C warmer, and 5°C warmer conditions. From top to bottom: Buff-tailed bumblebee (*Bombus (Bombus) terrestris*, used a proxy for the Rusty patched bumblebee (*Bombus (Bombus) affinis*) and the yellow-banded bumblebee (*Bombus (Bombus) terricola*), freeze-avoidant pollinators; Diamondback moth (*Plutella xylostella*), freeze-avoidant pest; Canadian tiger swallowtail (*Papilio canadensis*), freeze-avoidant pollinator; Eastern tiger swallowtail (*Papilio glaucus*), freeze-avoidant pollinator; Woolly bear caterpillar (*Pyrrharctia isabella*), freeze-tolerant pollinator; Bean leaf beetle (*Ceratoma trifurcata*), freeze-tolerant pest; and Hoverfly (*Syrphus ribesii*), freeze-tolerant pollinator. Images of insects adapted from: *Rusty-patched bumblebee queen* by Miklasevskaja, M., 1971, https://val.vtecostudies.org/projects/vtbees/bombus-affinis/ Copyright 2024 by Vermont Center for Ecostudies; *Diamondback moth* 2006, https://en.wikipedia.org/wiki/Diamondback_moth; *Papilio canadensis* by Mdf, 2008, https://en.wikipedia.org/wiki/Papilio_canadensis; *Mosaic Gynandromorphs, Eastern Tiger Swallowtail (Papilio glaucus)* by Grace, K., 1979, https://www.floridamuseum.ufl.edu/100-years/object/eastern-tiger-swallowtail/ Copyright 2024 by Florida Museum of Natural History; *Pyrrharctia isabella* by Reago, A. and McClarren C., 2014 https://en.wikipedia.org/wiki/Pyrrharctia_isabella; *Adult bean leaf beetle* by University of Nebraska-Lincoln, 2024, https://cropwatch.unl.edu/soybean-management/insects-bean-leaf-beetle Copyright 1869-2024 by University of Nebraska-Lincoln; *Syrphus ribesii* by Aiwok, 2010, https://en.wikipedia.org/wiki/Syrphus_ribesii.

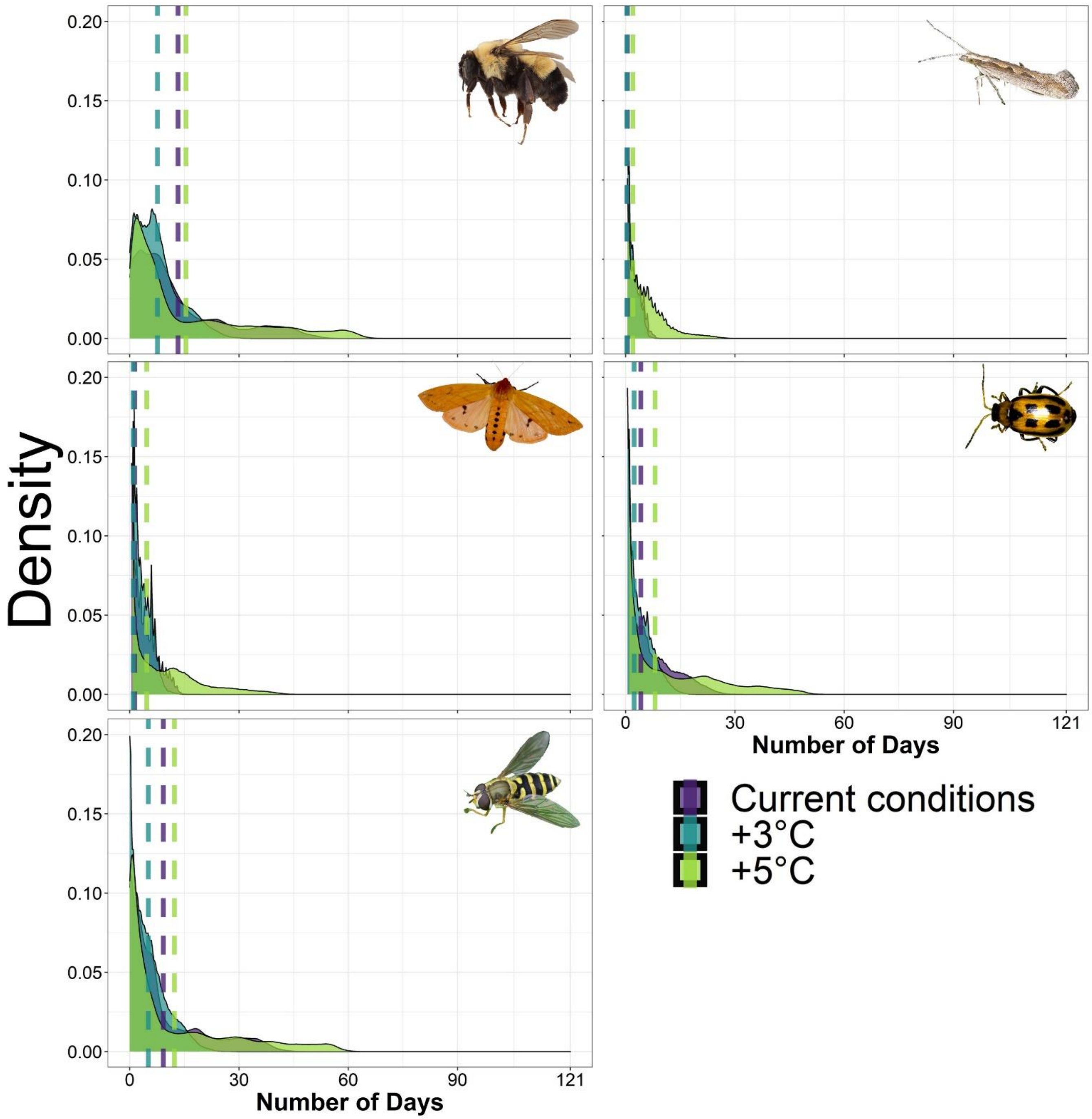


Figure S9. Density plots showing the distribution of total number of days in the winter season (December 1, 2016–March 31, 2017) below the lowest published supercooling point (i.e., best-case scenario) for insect species differing in their cold tolerance strategies and ecosystem services under current, 3°C warmer, and 5°C warmer conditions. Top row: Buff-tailed bumblebee (*Bombus (Bombus)*

*terrestris*, used a proxy for the Rusty patched bumblebee (*Bombus (Bombus) affinis*) and the yellow-banded bumblebee (*Bombus (Bombus) terricola*), freeze-avoidant pollinators, and the Diamondback moth (*Plutella xylostella*), freeze-avoidant pest. Middle row: Woolly bear caterpillar (*Pyrrharctia isabella*), freeze-tolerant pollinator and Bean leaf beetle (*Ceratoma trifurcata*), freeze-tolerant pest. Bottom row: Hoverfly (*Syrphus ribesii*), freeze-tolerant pollinator.  The two butterfly species (Canadian tiger swallowtail (*Papilio canadensis*) and Eastern tiger swallowtail (*Papilio glaucus*, both freeze-avoidant pollinators) are not pictured because under each climate scenario the total days below each of their SCPs was zero. Images of insects adapted from: *Rusty-patched bumblebee queen* by Miklasevskaja, M., 1971, https://val.vtecostudies.org/projects/vtbees/bombus-affinis/ Copyright 2024 by Vermont Center for Ecostudies; *Diamondback moth* 2006, https://en.wikipedia.org/wiki/Diamondback_moth; *Pyrrharctia isabella* by Reago, A. and McClarren C., 2014 https://en.wikipedia.org/wiki/Pyrrharctia_isabella; *Adult bean leaf beetle* by University of Nebraska-Lincoln, 2024,  https://cropwatch.unl.edu/soybean-management/insects-bean-leaf-beetle Copyright 1869-2024 by University of Nebraska-Lincoln; *Syrphus ribesii* by Aiwok, 2010, https://en.wikipedia.org/wiki/Syrphus_ribesii.

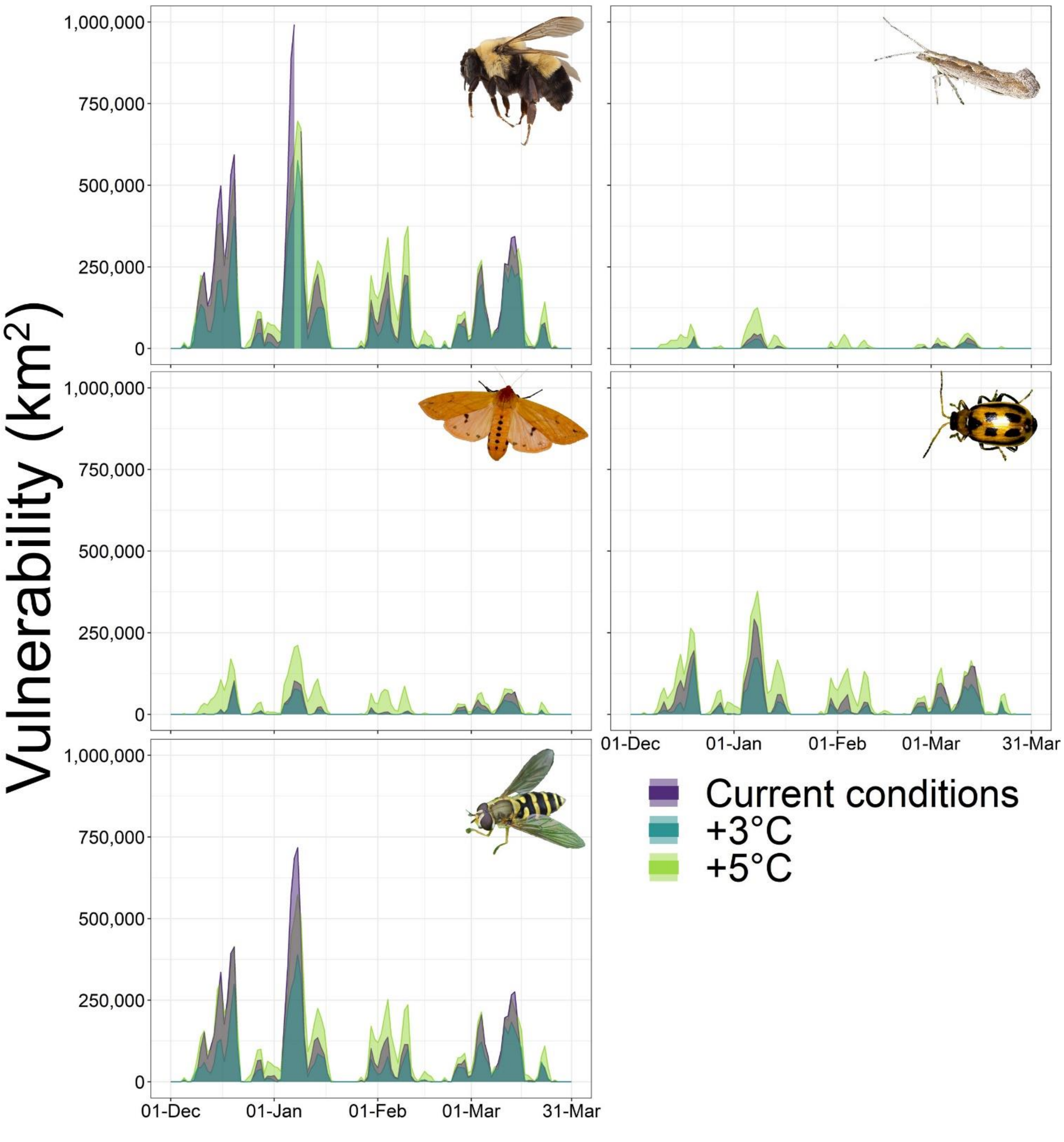


Figure S10. Daily extent of vulnerability (i.e., square kilometers where predicted ground temperatures were below the lowest published supercooling point (i.e. best-case scenario)) for insect species differing in their cold tolerance strategies and ecosystem services under current, 3°C warmer, and 5°C warmer conditions. Top row: Buff-tailed bumblebee (*Bombus (Bombus) terrestris*, used a proxy for the

Rusty patched bumblebee (*Bombus (Bombus) affinis*) and the yellow-banded bumblebee (*Bombus (Bombus) terricola*), freeze-avoidant pollinators, and the Diamondback moth (*Plutella xylostella*), freeze-avoidant pest. Middle row: Woolly bear caterpillar (*Pyrrharctia isabella*), freeze-tolerant pollinator and Bean leaf beetle (*Ceratoma trifurcata*), freeze-tolerant pest. Bottom row: Hoverfly (*Syrphus ribesii*), freeze-tolerant pollinator. The two butterfly species (Canadian tiger swallowtail (*Papilio canadensis*) and Eastern tiger swallowtail (*Papilio glaucus*, both freeze-avoidant pollinators) are not pictured because under each climate scenario the total days below each of their SCPs was zero. Images of insects adapted from: *Rusty-patched bumblebee queen* by Miklasevskaja, M., 1971, https://val.vtecostudies.org/projects/vtbees/bombus-affinis/ Copyright 2024 by Vermont Center for Ecostudies; *Diamondback moth* 2006, https://en.wikipedia.org/wiki/Diamondback_moth; *Pyrrharctia isabella* by Reago, A. and McClarren C., 2014 https://en.wikipedia.org/wiki/Pyrrharctia_isabella; *Adult bean leaf beetle* by University of Nebraska-Lincoln, 2024, https://cropwatch.unl.edu/soybean-management/insects-bean-leaf-beetle Copyright 1869-2024 by University of Nebraska-Lincoln; *Syrphus ribesii* by Aiwok, 2010, https://en.wikipedia.org/wiki/Syrphus_ribesii.

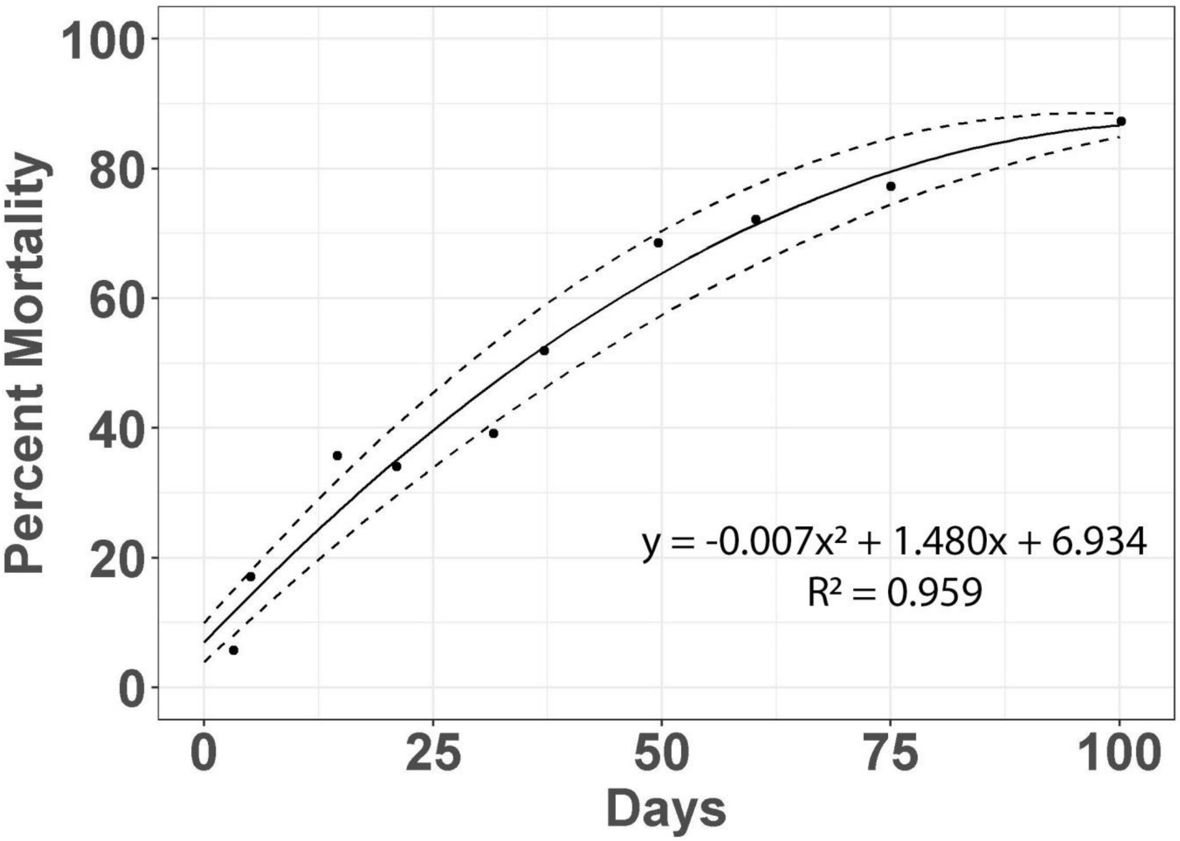


Figure S11. Linear regression model created using data points extracted from Lam and Pedigo (2000) on percent mortality as a function of time (n=30) in the bean leaf beetle (*Ceratoma trifurcata)*. From this regression model we extracted the fitted value ± confidence interval for 50% mortality.

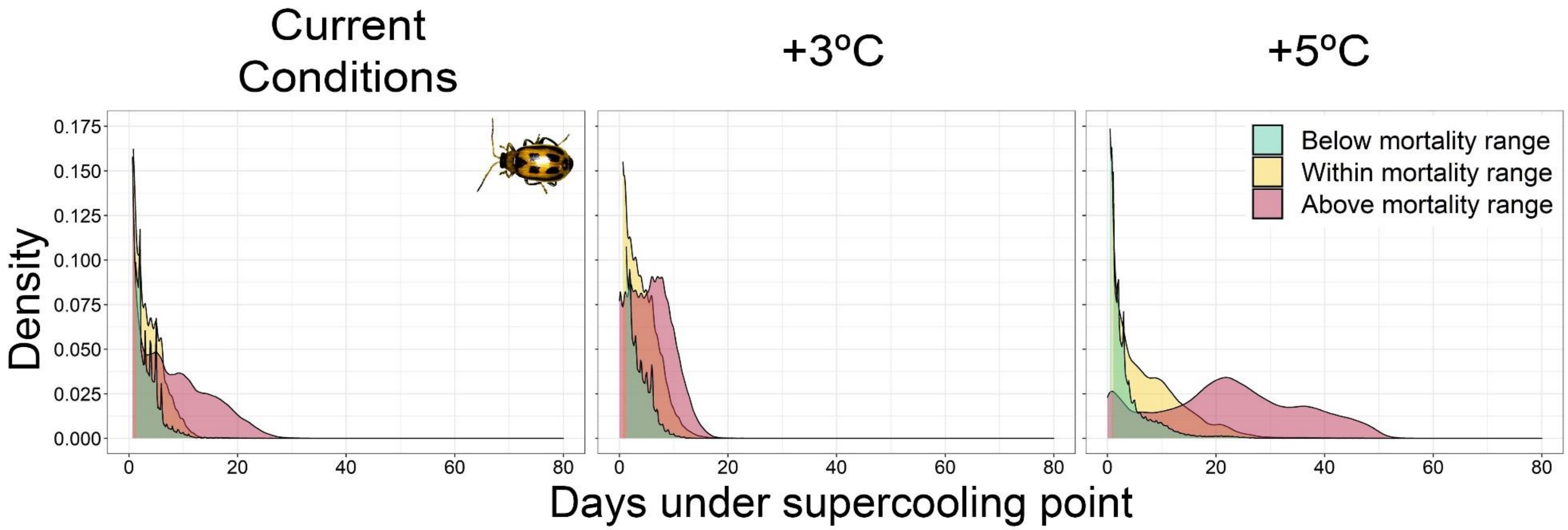


Figure S12. For three warming scenarios: current conditions (i.e. no warming), warming of 3°C, and warming of 5°C, density plots showing the overlap between the number of days during the winter season (December 1, 2016–March 31, 2017) with ground temperatures under the lowest published supercooling point (-9.3°C, i.e., best-case scenario) for bean leaf beetles (*C. trifurcata*) and the duration of consecutive days with ground temperatures at or below 0°C that would lead to 50% mortality based on experimental data extracted from Lam and Pedigo (2000). Lam and Pedigo (2000) found that the mean number of days until 50% mortality in a sample of *C. trifurcata* held at constant temperatures of 0°C was 34.6 days [28.9, 41.3]. ‘Above mortality range’ corresponds to consecutive sub-0°C days above this upper confidence limit of what *C. trifurcata* can withstand (i.e., least favorable scenario), ‘Within mortality range’ corresponds to consecutive sub-0°C days within the reported range for 50% mortality (i.e., moderate scenario), and

‘Below mortality range’ corresponds to sub-0°C days below the lower confidence limit of what *C. trifurcata* can withstand (i.e., most favorable scenario). Image of insect adapted from *Adult bean leaf beetle* by University of Nebraska-Lincoln, 2024, https://cropwatch.unl.edu/soybean-management/insects-bean-leaf-beetle Copyright 1869-2024 by University of Nebraska-Lincoln.

Supplementary References


Alford, D. V. 1969. A study of hibernation of bumblebees (Hymenoptera-Bombidae) in southern England. - Journal of Animal Ecology 38: 149-170.

Alford, D. V. 1975. Bumblebees. - Davis-Poynter.

Berzitis, E. A., et al. 2017. Winter warming effects on overwinter survival, energy use, and spring emergence of Cerotoma trifurcata (Coleoptera: Chrysomelidae). - Agricultural and Forest Entomology 19: 163-170.

Brown, C. L., et al. 2004. Freezing induces a loss of freeze tolerance in an overwintering insect. - Proceedings of the Royal Society B-Biological Sciences 271: 1507-1511.

Cameron, S. A., et al. 2007. A comprehensive phylogeny of the bumble bees (Bombus). - Biological Journal of the Linnean Society 91: 161-188.

Carrillo, M. A., et al. 2005. Supercooling point of bean leaf beetle (Coleoptera : Chrysomelidae) in Minnesota and a revised predictive model for survival at low temperatures. - Environmental Entomology 34: 1395-1401.

Carroll, T., et al. 2001. NOHRSC Operations and the Simulation of Snow Cover Properties for the Conterminous U.S. - In: Proceedings of the 69th Annual Meeting of the Western Snow Conference.

Dancau, T., et al. 2018. Elusively overwintering: a review of diamondback moth (Lepidoptera: Plutellidae) cold tolerance and overwintering strategy. - Canadian Entomologist 150: 156-173.

Dosdall, L. M., et al. 2011. The diamondback moth in canola and mustard: current pest status and future prospects. - Prairie Soils and Crops 4: 66-76.

Furlong, M. J., et al. 2013. Diamondback Moth Ecology and Management: Problems, Progress, and Prospects. - In: M. R. Berenbaum (ed) Annual Review of Entomology, Vol 58. pp. 517-541.

Goettel, M. S. and Philogene, B. J. R. 1987. Effects of photoperiod and temperature on the development of a univoltine population of the banded woollybear, *Pyrrharctia (Isia) isabella*. - Journal of Insect Physiology 24: 523-527.

Goulson, D. 2010. Bumblebees: Behaviour, Ecology, and Conservation. - Oxford University Press, Inc.

Grixti, J. C., et al. 2009. Decline of bumble bees (Bombus) in the North American Midwest. - Biological Conservation 142: 75-84.

Gu, H. N. 2009. Cold Tolerance and Overwintering of the Diamondback Moth (Lepidoptera: Plutellidae) in Southeastern Australia. - Environmental Entomology 38: 524-529.

Hagen, R. H. and Scriber, J. M. 1989. Sex-linked diapause, color, and allozyme loci in *Papilio glaucus*-linkage analysis and significance in a hybrid zone. - Journal of Heredity 80: 179-185.

Harcourt, D. G. 1954. The biology and ecology of the diamondback moth, *Plutella maculipennis* Curtis, in eastern Ontario. - In: Cornell University.

Hart, A. J. and Bale, J. S. 1998. Factors affecting the freeze tolerance of the hoverfly Syrphus ribesii (Diptera : Syrphidae). - Journal of Insect Physiology 44: 21-29.

Heinrich, B. 1972. Physiology of brood incubation in the bumblebee queen, *Bombus vosnesenskii*. - Nature 239: 223-225.

Heinrich, B. 1993. The Hot-Blooded Insects. - Springer-Verlag.

Homer, C. G., et al. 2012. The National Land Cover Database. - In: U.S. Geological Survey Fact Sheet.

Hunt, T. E., et al. 1995. Bean leaf beetle injury to seedling soybean - consumption, effects of leaf expansion, and economic injury levels. - Agronomy Journal 87: 183-188.

Idris, A. B. and Grafius, E. J. 1996. Evidence of pre-imaginal overwintering of diamondback moth, Plutella xylostella (Lepidoptera: Plutellidae) in Michigan. - Great Lakes Entomologist 29: 25-30.

Kimura, T., et al. 1987. Overwintering of the diamondback moth, *Plutella xylostella* Linne, in Aomoria Prefecture. 3. Mortality of the sectond instar larvae under the snow. - Annual Report of the Society of Plant Protection of North Japan 38: 141-142.

Kimura, T. and Fujimura, T. 1988. Overwintering of the diamondback moth, *Plutella xylostella* Linne, in Aomori Prefecture 4. Longevity of eggs, larvae, papae and adults under the snow. - Annual Report of the Society of Plant Protection of North Japan 39: 229-231.

Kukal, O., et al. 1991. Cold tolerance of the pupae in relation to the distribution of swallowtail butterflies. - Canadian Journal of Zoology-Revue Canadienne De Zoologie 69: 3028-3037.

Lam, W. K. F. and Pedigo, L. P. 2000. Cold tolerance of overwintering bean leaf beetles (Coleoptera : Chrysomelidae). - Environmental Entomology 29: 157-163.

Larrere, M., et al. 1993. Juvenile-hormone biosynthesis and diapause termination in *Bombus terrestris*. - Invertebrate Reproduction & Development 23: 7-14.

Layne, J. R., et al. 1999. Cold hardiness of the woolly bear caterpillar (Pyrrharctia isabella Lepidoptera : Arctiidae). - American Midland Naturalist 141: 293-304.

Liu, S. S., et al. 2002. Development and survival of the diamondback moth (Lepidoptera : Plutellidae) at constant and alternating temperatures. - Environmental Entomology 31: 221-231.

Loughran, J. C. and Ragsdale, D. W. 1986. Life-cycle of the bean leaf beetle, *Ceratoma trifurcata* (Coleoptera, Chrisomelidae), in southern Minnesota. - Annals of the Entomological Society of America 79: 34-38.

Marshall, K. E. and Sinclair, B. J. 2011. The sub-lethal effects of repeated freezing in the woolly bear caterpillar Pyrrharctia isabella. - Journal of Experimental Biology 214: 1205-1212.

McCreary, C. 2013. Development of a degree day model and economica thresholds for *Ceratoma trifurcata* (Coleoptera: Chrysomelidae). - In: University of Guelph.

Mercader, R. J. and Scriber, J. M. 2008. Asymmetrical thermal constraints on the parapatric species boundaries of two widespread generalist butterflies. - Ecological Entomology 33: 537-545.

Mesinger, F., et al. 2006. North American Regional Reanalysis. - Bulletin of the American Meteorological Society 87: 343-360.

Moraiti, C. A. and Papadopoulos, N. T. 2017. Obligate annual and successive facultative diapause establish a bet-hedging strategy of Rhagoletis cerasi (Diptera: Tephritidae) in seasonally unpredictable environments. - Physiological Entomology 42: 225-231.

National Hydrologic Remote Sensing Center 2004. Snow Data Assimilation System (SNODAS) Data Products at NSIDC, Version 1. - In: National Snow and Ice Data Center.

Owen, E. L., et al. 2013. Can Winter-Active Bumblebees Survive the Cold? Assessing the Cold Tolerance of Bombus terrestris audax and the Effects of Pollen Feeding. - Plos One 8:

Park, Y. and Kim, Y. 2014. A specific glycerol kinase induces rapid cold hardening of the diamondback moth, Plutella xylostella. - Journal of Insect Physiology 67: 56-63.

Pedigo, L. P. 1994. Bean leaf beetle. - In: L. G. Higley and D. J. Boethel (eds), Handbook of Soybean Insect Pests. Entomological Society of America, pp. 42-44.

Razumov, V. P. 1970. How the diamond-back moth overwinters. - Zashchita Rastenii 15:

Rockey, S. J., et al. 1987. A latitudinal and obligatory diapause response in three subspecies of the eastern tiger swallowtail *Papilio glaucus* (Lepidoptera: Papilionidae). - American Midland Naturalist 118: 162-168.

Ryan, S. F., et al. 2018. The role of latitudinal, genetic and temperature variation in the induction of diapause of Papilio glaucus (Lepidoptera: Papilionidae). - Insect Science 25: 328-336.

Saito, O. 1994. Tolerance of the diamondback moth, *Plutella xylostella* (L.) (Lepidoptera: Yponomeutidaie), to low constant temperature. - Annual Report of the Society of Plant Protection of North Japan 45: 158-159.

Scriber, J. M. 2011. Impacts of climate warming on hybrid zone movement: Geographically diffuse and biologically porous "species borders". - Insect Science 18: 121-159.

Sgolastra, F., et al. 2010. Effect of temperature regime on diapause intensity in an adult-wintering Hymenopteran with obligate diapause. - Journal of Insect Physiology 56: 185-194.

Thornton, P. E., et al. 2016. Daymet: Daily Surface Weather Data on a 1-km Grid for North America, Version 3. - In: O. DAAC (ed).

Williams, C. M., et al. 2012. Lepidopteran species differ in susceptibility to winter warming. - Climate Research 53: 119-130.

Zhang, L. J., et al. 2015. Identification of heat shock protein genes hsp70s and hsc70 and their associated mRNA expression under heat stress in insecticide-resistant and susceptible diamondback moth, Plutella xylostella (Lepidoptera: Plutellidae). - European Journal of Entomology 112: 215-226.
